# Mitochondrial dynamics regulate genome stability via control of caspase-dependent DNA damage

**DOI:** 10.1101/2021.09.13.460067

**Authors:** Kai Cao, Joel S Riley, Catherine Cloix, Yassmin Elmasry, Gabriel Ichim, Kirsteen J Campbell, Stephen WG Tait

## Abstract

Mitochondrial dysfunction is interconnected with cancer. Nevertheless, how defective mitochondria promote cancer is poorly understood. We find that mitochondrial dysfunction promotes DNA damage under conditions of increased apoptotic priming. Underlying this process, we reveal a key role for mitochondrial dynamics in the regulation of DNA damage and genome instability. The ability of mitochondrial dynamics to regulate oncogenic DNA damage centres upon the control of minority MOMP, a process that enables non-lethal caspase activation leading to DNA damage. Mitochondrial fusion suppresses minority MOMP, and its associated DNA damage, by enabling homogenous mitochondrial expression of anti-apoptotic BCL-2 proteins. Finally, we find that mitochondrial dysfunction inhibits pro-apoptotic BAX retrotranslocation, causing BAX mitochondrial localization thereby promoting minority MOMP. Unexpectedly, these data reveal oncogenic effects of mitochondrial dysfunction that are mediated via mitochondrial dynamics and caspase-dependent DNA damage.

## Introduction

Mitochondrial dysfunction has pleiotropic impact on cancer (Giampazolias and Tait, 2016). For instance, mitochondrial respiratory complex proteins and TCA enzymes bearing tumour associated mutations, generate oncometabolites (Isaacs et al., 2005; Pollard et al., 2007; Sciacovelli et al., 2016; Selak et al., 2005). Moreover, loss of function mutations in mitochondrial DNA (mtDNA) are common in cancer and have been shown to accelerate tumorigenesis (Gorelick et al., 2021; Smith et al., 2020). Nonetheless, how dysfunctional mitochondria promote cancer remains largely an open question.

While inhibition of mitochondrial apoptosis has well established oncogenic effects, through increased apoptotic priming, tumour cells are often sensitized to cell killing cancer therapies (Certo et al., 2006; Singh et al., 2019). Mitochondria regulate apoptosis via mitochondrial outer membrane permeabilization or MOMP (Bock and Tait, 2020). This key event releases soluble mitochondrial intermembrane space proteins into the cytoplasm, notably cytochrome *c*, that activate caspases proteases causing rapid cellular demise. Because it dictates cell fate, mitochondrial outer membrane integrity is tightly regulated by BCL-2 protein family members (Campbell and Tait, 2018).

MOMP is usually considered a lethal point-of-no-return due to its extensive nature, often occurring in all mitochondria, coupled to an invariable loss of mitochondrial function (Goldstein et al., 2000; Lartigue et al., 2009; Rehm et al., 2003). However, we have previously described conditions whereby MOMP can be heterogenous permitting cell survival (Ichim et al., 2015; Tait et al., 2010). Following a sub-lethal stress, a limited mitochondrial cohort selectively permeabilizes, which we termed minority MOMP (Ichim et al., 2015). Strikingly, minority MOMP can engage sub-lethal caspase activity promoting DNA damage that is dependent upon caspase-activated DNAse (CAD) (Ichim et al., 2015). By causing DNA damage, minority MOMP may contribute to the paradoxical oncogenic effects of apoptotic signaling reported in different studies (Ichim and Tait, 2016). Moreover, minority MOMP has been recently implicated in an expanding array of functions including increased cancer aggressiveness, innate immunity and inflammation triggered by mtDNA double-strand breaks (Berthenet et al., 2020; Brokatzky et al., 2019; Tigano et al., 2021).

Here, we investigated the relationship between mitochondrial dysfunction and DNA damage. Surprisingly, we uncovered a key role for mitochondrial dynamics in the regulation of DNA damage. Mitochondrial fission, a consequence of mitochondrial dysfunction, promotes minority MOMP causing caspase-dependent DNA damage and genome instability. Secondly, we find reduced retrotranslocation of pro-apoptotic BAX on dysfunctional mitochondria, thus facilitating minority MOMP. These data reveal an unanticipated link between mitochondrial dysfunction and oncogenic DNA damage that is mediated through minority MOMP and caspase activity.

## Results

### Mitochondrial dynamics regulate DNA damage

We aimed to understand how mitochondrial dysfunction can be oncogenic. Given the tumor promoting roles of DNA damage, we initially investigated its interconnection with mitochondrial function. To cause mitochondrial dysfunction, U2OS and HeLa cells were treated with the uncoupler carbonyl cyanide *m*-chlorophenyl hydrazone (CCCP). In order to phenocopy increased apoptotic priming that is found in pre-malignant and tumour cells, we co-treated cells with ABT-737, a BH3-mimetic compound that selectively neutralises anti-apoptotic BCL-2, BCL-xL and BCL-w. The response to DNA damage was measured by ɣH2AX staining and flow cytometry. In both HeLa and U2OS cells BH3-mimetic treatment led to an increase in ɣH2AX positive cells that was significantly enhanced by combined treatment with CCCP, consistent with mitochondrial dysfunction promoting DNA damage (**Figure 1A**). Mitochondrial dynamics and function are tightly interconnected such that mitochondrial dysfunction causes mitochondrial fission (Oltersdorf et al., 2005). We therefore investigated whether mitochondrial dynamics affected DNA damage triggered by BH3-mimetic treatment. To disrupt mitochondrial fusion, we used *Mfn1/2*^-/-^ murine embryonic fibroblasts (*Mfn1/2*^-/-^ MEF) and, as control, reconstituted these cells with MFN2 (*Mfn1*^-/-^ MEF). As expected, *Mfn1/2*^-/-^ MEF displayed a hyper-fragmented mitochondrial network whereas MFN2 reconstitution of these cells (*Mfn1*^-/-^) restored mitochondrial fusion, resulting in a filamentous mitochondrial network (**Figure 1B, C**). *Mfn1/2*^-/-^ and *Mfn1*^-/-^ MEF were treated with ABT-737 (10 µM, 3 hours) and the DNA damage response was assessed by analyzing γH2AX levels by western blot or by flow cytometry (**Figures 1D, E**). *Mfn1/2*^-/-^ MEF exhibited increased γH2AX, consistent with mitochondrial fission promoting DNA damage. Because DNA damage can be oncogenic, we investigated if cells with extensive mitochondrial fission were more prone to transformation. *Mfn1/2^-/-^* and *Mfn1^-/-^* MEF were passaged repeatedly in ABT-737. Following treatment, cells were assayed for transformation *in vitro* by determining anchorage-independent growth in soft agar. Specifically following culture in ABT-737, *Mfn1/2^-/-^* MEF formed colonies more readily *Mfn1^-/-^* MEF (**Figure 1F, G**). In reciprocal fashion, we investigated the impact of inhibiting mitochondrial fission upon DNA damage. DRP1 plays a central role in mitochondrial fission (Ishihara et al., 2009; Wakabayashi et al., 2009). To inhibit mitochondrial fission we used *Drp1^fl/fl^* MEF, which when infected with adenoviral Cre efficiently delete DRP1, causing a hyper-fused mitochondrial network (**Figure 1H, Supplementary Figure 1A***). Drp1^fl/fl^* and *Drp1^-/-^* MEF were treated with ABT-737 and γH2AX was measured by flow cytometry, as before. MEF expressing DRP1 have elevated levels of γH2AX after exposure to ABT-737, but this was completely abolished in DRP1-deficient cells (**Figure 1I**). These data show that mitochondrial dysfunction and fission promote oncogenic DNA damage and transformation.

**Figure 1.**
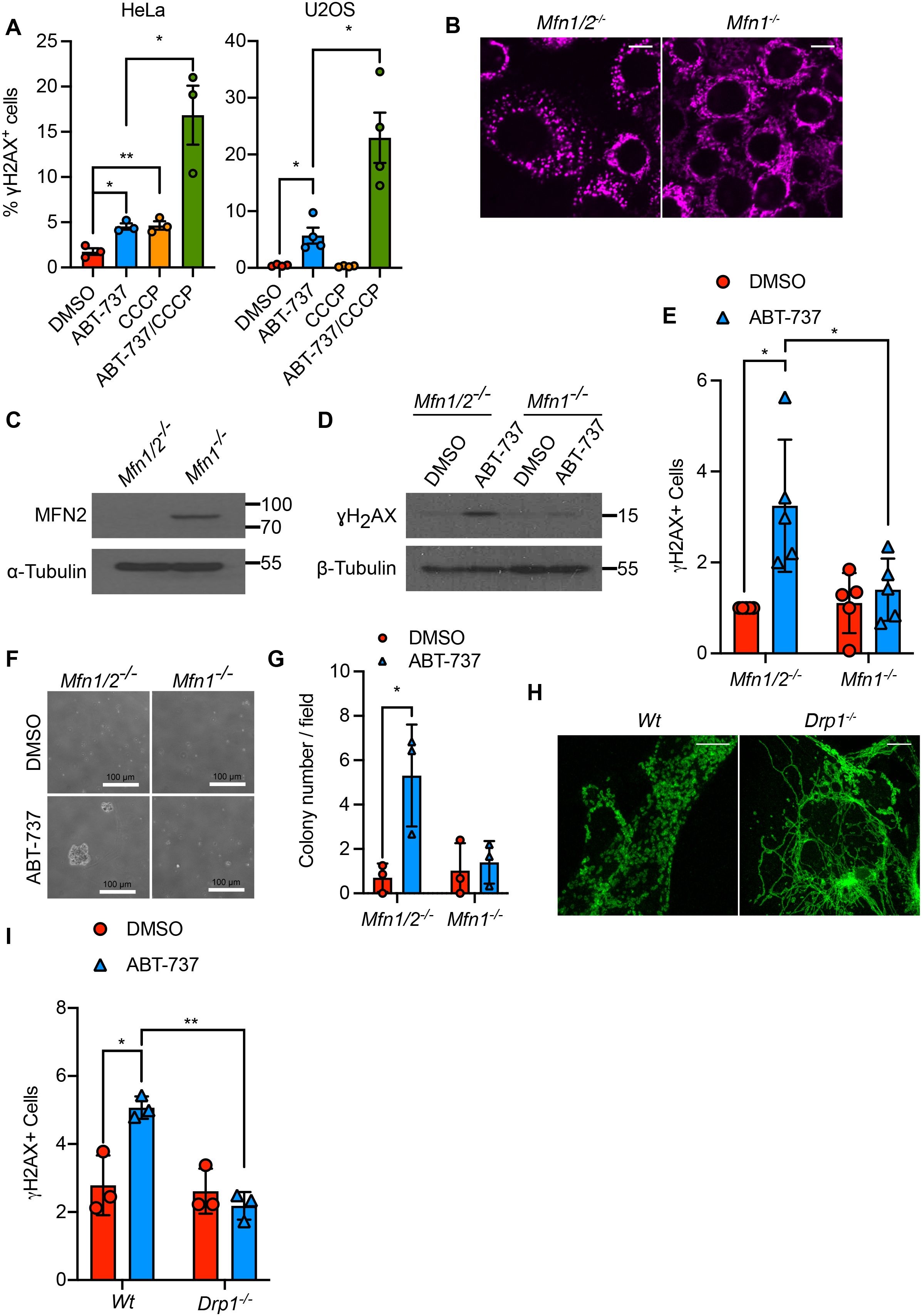
Mitochondrial dynamics regulate DNA damage. A) Flow cytometric analysis of HeLa and U2OS cells treated with 10 µM CCCP for 30 min before treatment with 10 µM ABT-737 for 3 h. Cells were immunostained with anti-ɣH2AX antibody. Data represented as mean ± SEM from 3 independent experiments. B) Airyscan images of *Mfn1/2*^-/-^ and *Mfn1*^-/-^ MEF, immunostained with anti-TOM20 antibody. Scale bar = 10 µm. C) Immunoblot of MFN2 in *Mfn1/2*^-/-^ and *Mfn1*^-/-^ MEF. D) ɣH2AX expression in *Mfn1/2*^-/-^ and *Mfn1*^-/-^ MEF treated with 10 µM ABT-737 for 3 h. E) Flow cytometric analysis of ɣH2AX expression in *Mfn1/2*^-/-^ and *Mfn1*^-/-^ MEF treated with 10 µM ABT-737 for 3 h. Data represented as mean ± SEM from 5 independent experiments. F) *Mfn1/2*^-/-^ and *Mfn1*^-/-^ MEF were cultured for twenty passages in 10 µM ABT-737 and their anchorage-independent growth assessed by soft agar assay. Representative images from 3 independent experiments shown. G) Quantification of anchorage-independent growth in soft agar from (D). Data are expressed as mean ± SD from 3 independent experiments and analysed using student’s t-test. H) Airyscan images of *Drp1^fl/fl^* MEF infected with AdCre and immunostained with anti-TOM20 antibody. Scale bar = 10 µm. I) Flow cytometric analysis of ɣH2AX expression in Wt and *Drp1^fl/fl^* MEF treated with 10 µM ABT-737 for 3 h. Data are expressed at mean ± SEM from 3 independent experiments and analysed using student’s t-test.

### Mitochondrial dynamics regulate DNA damage and genome-instability in a caspase and CAD dependent manner

We next sought to understand how mitochondrial dynamics regulate DNA damage. Because we had found that pro-apoptotic BH3-mimetic treatment potentiated DNA damage, we investigated a role for apoptotic caspase function. Wild type MEF or MEF overexpressing DRP1 were treated with the pan-caspase inhibitor qVD-OPh and γH2AX was measured by flow cytometry (**Figure 2A, B, Supplementary Figure 2A, B**). MEF cells overexpressing DRP1 displayed a more fragmented mitochondrial network and had higher levels of γH2AX compared to their empty vector counterparts, this is consistent with our earlier data. Crucially, γH2AX was prevented by treatment with the pan-caspase inhibitor qVD-OPh, demonstrating a key role for caspase activation in DNA damage (**Figure 2B**). Given these findings, we investigated a possible correlation between expression of the mitochondrial fission protein DRP1 and mutational burden in cancer. TCGA PanCancer Atlas studies were investigated through cBioportal. Of these, a significant association between increased mutational count in *DNM1L* mRNA high quartile versus *DNM1L* mRNA low quartile was found in invasive breast carcinoma and lung adenocarcinoma (out of 22 studies) with the inverse relationship not observed in any cancer type (**Figures 2C, 2D, Supplemental Table 1** and data not shown). In both invasive breast cancer and lung adenocarcinoma, DNA damage response pathways were enriched in the *DNM1L* mRNA high quartile consistent with engagement of DNA damage (**Supplementary Figures 2C, D, E**). To further investigate the role of caspase activity, we investigated the impact of mitochondrial dynamics upon genome instability. To this end, we used the PALA assay, in which gene amplification of CAD (carbamyl phosphate synthetase/aspartate transcarbamylase/dihydro-orotase, note that this is distinct from caspase-activated DNase described later) enables resistance to PALA (N-phosphonoacetyl-L-aspartate)(Wahl et al., 1979). To determine if alterations in mitochondrial dynamics, also affect genome instability dependent upon caspase activity, we passaged *Mfn1/2^-/-^* and *Mfn1^-/-^* MEF with sub-lethal doses of ABT-737 in the presence or absence of QVD-OPh. Following treatment, cells were grown in the presence of PALA and clonogenic survival was measured (**Figure 2E**). Importantly, ABT-737 treated *Mfn1/2^-/-^* MEF gave significantly more colonies than *Mfn1^-/-^* following PALA treatment, in a caspase-dependent manner (**Figure 2E, F**). In line with increased survival following PALA treatment, qPCR revealed amplification of the *Cad* locus only in *Mfn1/2^-/-^* MEF repeatedly treated with ABT-737 (**Figure 2G**). We and others have previously found that non-lethal caspase activity can cause DNA damage and genome instability dependent upon caspase-activated DNAse (CAD) (Ichim et al., 2015; Lovric and Hawkins, 2010). To examine the role of CAD in genomic instability we used the *Mfn1/2^-/-^* and *Mfn1^-/-^* MEF in which we deleted the *Dff40* gene (encoding CAD) using CRISPR-Cas9 genome editing (**Supplementary Figure 2F**). As before, *Mfn1/2^-/-^* cells resisted PALA treatment and efficiently grew as colonies following ABT-737 treatment, whereas *Mfn1^-/-^* cells did not (**Figure 2H, I**). However, deletion of CAD completely abrogated clonogenic potential. *Cad* DNA expression and anchorage-independent growth were also diminished following ABT-737 treatment in CAD/*Dff40* deleted cells as compared to their controls (**Figure 2J, K, Supplementary Figure 2G**). Together, these data show that mitochondrial fission promotes genome instability in a caspase and CAD dependent manner.

**Figure 2.**
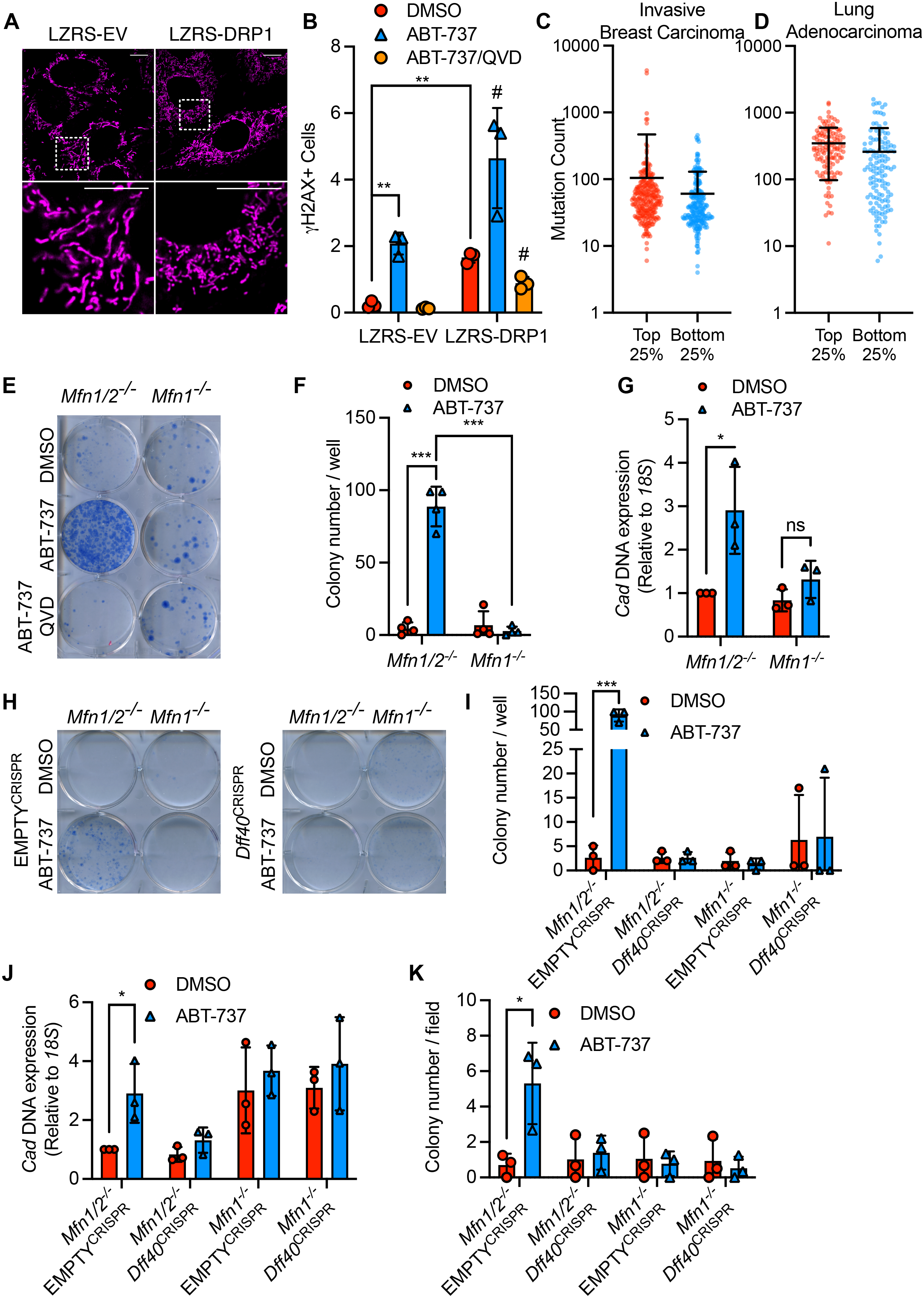
Mitochondrial dynamics regulate DNA damage and genome-instability in a caspase and CAD dependent manner. A) Airyscan images of MEF overexpressing LZRS-DRP1 or LZRS empty vector, stained with MitoTracker Deep Red. Scale bar = 10 µm. B) Flow cytometric analysis of MEF stably over-expressing LZRS control or LZRS-DRP1, treated with 10 µM ABT-737 with and without 20 µM QVD for 3 h. Data are expressed at mean ± SEM from 3 independent experiments and analysed using student’s t-test. C) Mutation counts in patient lung adenocarcinoma samples from the highest and lowest DNM1L mRNA quartiles. Significance is analysed by Mann-Whitney test. Data points represent individual patient samples, bar represents mean ± SD. D) Mutation counts in patient breast invasive carcinoma cancer samples from the highest and lowest DNM1L mRNA quartiles. Significance is analysed by Mann-Whitney test. Data points represent individual patient samples, bar represents mean ± SD. E) *Mfn1/2*^-/-^ and *Mfn1*^-/-^ MEF were treated daily for twenty passages with 10 µM ABT-737 with and without 20 µM QVD. Clonogenic survival was performed in the presence of 100 µM PALA. Data is a representative example of 4 independent experiments. F) Quantification of clonogenic outgrowth from (A) from 4 independent experiments. Data are expressed as mean ± SD and analysed using student’s t-test. G) Quantification of Cad DNA levels in *Mfn1/2*^-/-^ and *Mfn1*^-/-^ MEF treated with or without 10 µM ABT-737. Data are expressed as mean ± SD from 3 independent experiments and analysed using student’s t-test. H) *Mfn1/2*^-/-^ and *Mfn1*^-/-^ MEF with and without CRISPR-Cas9-mediated *Dff40* deletion treated daily for twenty passages with 10 µM ABT-737 with and without 20 µM QVD. Clonogenic survival was performed in the presence of 100 µM PALA. Data is a representative example of 3 independent experiments. I) Quantification of clonogenic outgrowth from (F) from 3 independent experiments. Data are expressed as mean ± SD and analysed using student’s t-test. J) Quantification of *Cad* DNA levels in *Mfn1/2*^-/-^ and *Mfn1*^-/-^ MEF with and without *Dff40* deletion, and treated with or without 10 µM ABT-737. Data are expressed as mean ± SD from 3 independent experiments, and analysed using student’s t-test. K) *Mfn1/2*^-/-^ and *Mfn1*^-/-^ MEF with and without *Dff40* deletion cultured for twenty passages in 10 µM ABT-737 and their anchorage-independent growth assessed by soft agar assay. Data are expressed as mean ± SD from 3 independent experiments, and analysed using student’s t-test.

### Minority MOMP occurs on fragmented mitochondria and is regulated by mitochondrial dynamics

We have previously found that permeabilization of limited mitochondria – called minority MOMP – can engage non-lethal caspase activity causing CAD activation and DNA damage. This knowledge, coupled to our previous data, led us to investigate a role for mitochondrial dynamics in the regulation of minority MOMP. To address this, we combined super-resolution Airyscan confocal microscopy together with our fluorescent reporter that allows detection of minority MOMP (Ichim et al., 2015). This reporter comprises cytosolic FKBP-GFP (cytoGFP) and mitochondrial inner membrane targeted FRB-mCherry (mito-mCherry). Upon loss of mitochondrial outer membrane integrity, and in the presence of chemical heterodimeriser (AP21967), these two proteins bind one another, recruiting cytoGFP to the permeabilised mitochondria. HeLa or U2OS were treated with a non-lethal dose of BH3-mimetic ABT-737 (10 µM) for 3 hours. Consistent with our previous data, this treatment was sufficient to engage minority MOMP, as evidenced by localisation of cytoGFP to specific mitochondria (**Figure 3A**). Super-resolution analysis of these mitochondria revealed that selectively permeabilised mitochondria were separate from the mitochondria network, suggesting that minority MOMP preferentially occurs on fragmented mitochondria (**Figure 3A, B**). Extensive mitochondrial fission is a well-established consequence of MOMP (Bhola et al., 2009; Frank et al., 2001). Therefore, to determine whether mitochondria fragmentation was a cause or consequence of minority MOMP, U2OS cells expressing cytoGFP and mito-mCherry were imaged by live-cell microscopy. Treatment with ABT-737 (10 µM) led to minority MOMP, apparent by the translocation of cytoGFP into mitochondria after 124 minutes. Importantly, these mitochondria were fragmented from the mitochondrial network prior to cytoGFP translocation at 120 minutes (**Figure 3C**, **Movie 1**). This suggests that minority MOMP preferentially occurs on fragmented mitochondria. We next investigated a causal role for mitochondrial dynamics in regulating minority MOMP. We next used these cells to investigate a role for mitochondrial fusion in regulating minority MOMP. *Mfn1/2*^-/-^ and *Mfn1*^-/-^ MEF expressing the MOMP reporter, were treated with a sub-lethal dose of ABT-737. Strikingly, increased levels of minority MOMP were observed in *Mfn1/2*^-/-^ MEF when compared to *Mfn1*^-/-^ MEF (**Figure 3D**). This is consistent with minority MOMP occurring primarily on fragmented mitochondria, with mitochondrial fusion having an inhibitory effect. To further address this, we investigated the impact of inhibiting mitochondrial fission upon minority MOMP following treatment of *Drp1^fl/fl^* and *Drp1^-/-^* MEF with ABT-737. MEF expressing DRP1 undergo minority MOMP and after exposure to ABT-737, but this was completely abolished in DRP1-deleted cells (**Figure 3E**). Together, these data demonstrate that mitochondrial dynamics regulate minority MOMP; mitochondrial fusion is inhibitory whereas fission promotes minority MOMP.

**Figure 3.**
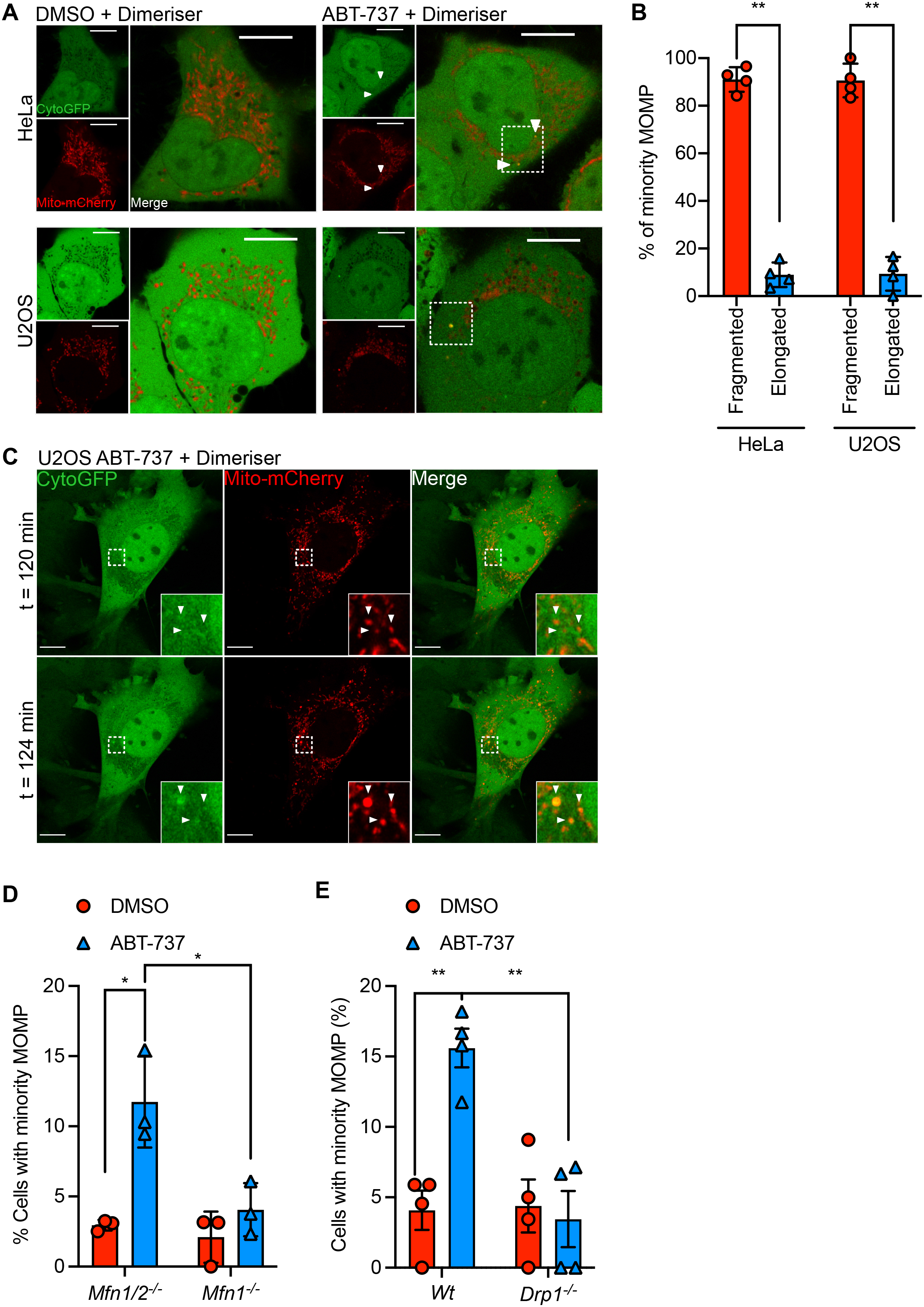
Minority MOMP occurs on fragmented mitochondria and is regulated by mitochondrial dynamics. A) Fixed super-resolution Airyscan images of HeLa and U2OS cells transfected with cyto-GFP (green) and mito-mCherry (red). Cells were treated with 10 µM ABT-737 for 3 h in the presence of dimeriser. Scale bar = 10 µm. B) Quantification of fragmentation or elongation of mitochondria which have undergone minority MOMP, as visualised in (A). Data represented as mean ± SD from 4 independent experiments and analysed using student’s t-test. C) Live-cell Airyscan imaging of U2OS cells transfected with cyto-GFP (green) and mito-mCherry (red) and treated with 10 µM ABT-737 in the presence of dimeriser. Scale bar = 10 µm. See **Movie 1**. D) Quantification of minority MOMP assessed in *Mfn1/2*^-/-^ and *Mfn1*^-/-^ MEF, transfected with cyto-GFP and mito-mCherry. Cells were treated with 10 µM ABT-737 for 3 h in the presence of dimeriser. Data represented as mean ± SEM from 3 independent experiments. E) Quantification of minority MOMP assessed in Wt and *Drp1^fl/fl^* MEF, transfected with cyto-GFP and mito-mCherry. Cells were treated with 10 µM ABT-737 for 3 h in the presence of dimeriser. Data represented as mean ± SEM from 4 independent experiments.

### Pro-survival BCL-2 proteins display inter-mitochondrial heterogeneity in expression

Our data demonstrate that mitochondrial fission promotes minority MOMP enabling caspase dependent DNA damage. Nevertheless, how mitochondrial dynamics regulate minority MOMP is not known. Mitochondrial outer membrane integrity is regulated by the balance of pro- and anti-apoptotic BCL-2 family proteins (Campbell and Tait, 2018). We hypothesised that inter-mitochondrial variation in BCL-2 family expression may underlie minority MOMP. To investigate this hypothesis, we aimed to visualise endogenous levels of BCL-2 family proteins on individual mitochondria. CRISPR-Cas9 genome editing can be used to knock-in fluorescent proteins at defined genomic loci to enable endogenous tagging of proteins (Bukhari and Muller, 2019). Using this approach, we generated clonal knock-in HeLa cell lines where the red fluorescent protein Scarlet was fused to the N-termini of BCL-2, BCL-xL and MCL-1. As verification, western blotting using antibodies specific BCL-2, BCL-xL, MCL-1 and Scarlet confirmed that these cell lines expressed these fusion proteins at similar levels to their endogenous counterparts (**Figure 4A, B**). Secondly, Airyscan super-resolution microscopy demonstrated mitochondrial localisation of Scarlet-BCL-2, BCL-xL and MCL-1, as expected (**Figure 4C**). Finally, we monitored cell viability using SYTOX Green exclusion and IncuCyte real-time imaging in response to BH3-mimetic treatment (ABT-737 and S63845). This demonstrated that all knock-in cell lines underwent cell death in response to BH3-mimetic treatment (**Supplementary Figure 3A**). Using these knock-in cells, we next acquired super-resolution microscopy images of Scarlet-tagged BCL-2, BCL-xL and MCL-1 then applied a colour grading lookup table (LUT) such that the brighter the Scarlet signal, the more magenta the image. This revealed heterogeneity of Scarlet BCL-2, BCL-xL and MCL-1 across the mitochondrial network (**Figure 4D**). Given our previous data, we hypothesised that BCL-2 family protein heterogeneity is regulated by mitochondrial dynamics. To test this, we inhibited mitochondrial fission through CRISPR-Cas9 deletion of DRP1. Western blot confirmed DRP1 deletion, resulting in extensive mitochondrial hyperfusion (**Supplementary Figure 3B, C**). Strikingly, cells with hyperfused mitochondria displayed much reduced inter-mitochondrial heterogeneity of MCL-1, BCL-2 or BCL-xL (**Figure 4E, F, Supplementary Figure 3D**). Combined, these data show that within a cell extensive inter-mitochondrial heterogeneity in BCL-2 expression exists that is profoundly impacted by mitochondrial dynamics.

**Figure 4.**
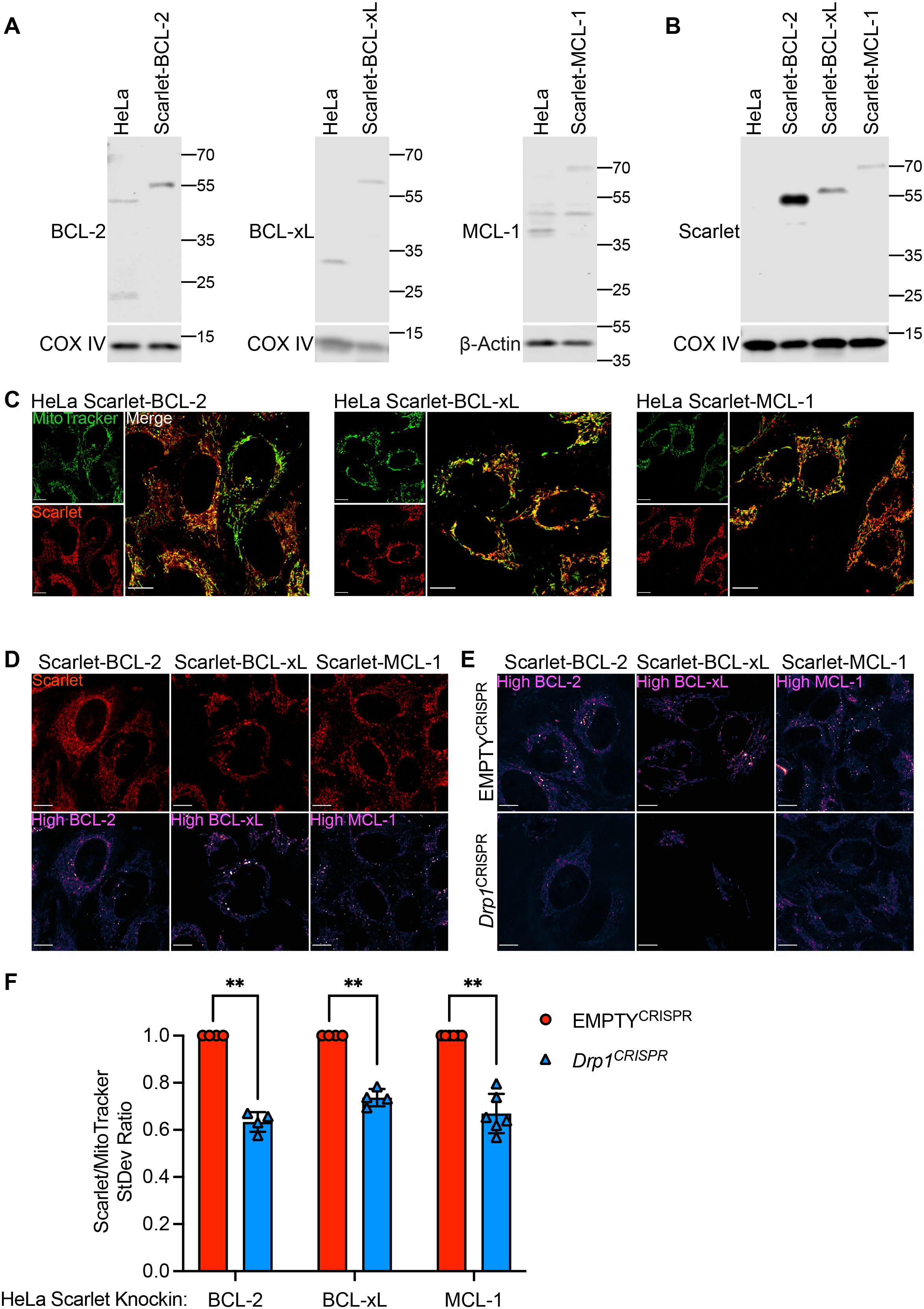
Pro-survival BCL-2 proteins display inter-mitochondrial heterogeneity in expression. A) Immunoblots of HeLa cells with CRISPR-Cas9-mediated knockin of Scarlet into the BCL-2, BCL-xL or MCL-1 locus using antibodies against BCL-2, BCL-xL or MCL-1. COX IV or β-actin serves as loading controls B) Immunoblots of HeLa cells with CRISPR-Cas9-mediated knockin of Scarlet into the BCL-2, BCL-xL or MCL-1 locus using an antibody against Scarlet. COX IV serves as a loading control. C) Live-cell Airyscan imaging of HeLa Scarlet-BCL-2, Scarlet-BCL-xL and Scarlet-MCL-1 cells. Cells were incubated with MitoTracker Green to stain mitochondria. D) Live-cell Airyscan imaging of HeLa Scarlet-BCL-2, Scarlet-BCL-xL and Scarlet-MCL-1 cells. Magenta LUT applied to reveal areas of high BCL-2, BCL-XL and MCL-1 expression. E) Live-cell Airyscan imaging of HeLa Scarlet-BCL-2, Scarlet-BCL-xL and Scarlet-MCL-1 cells with and without CRISPR-Cas9-mediated *Drp1* deletion. Magenta LUT applied to reveal areas of high BCL-2, BCL-XL and MCL-1 expression. F) Quantification of Scarlet to MitoTracker signal standard deviation in HeLa Scarlet-BCL-2, Scarlet-BCL-xL and Scarlet-MCL-1 cells, with and without CRISPR-Cas9-mediated *Drp1* deletion. Data are expressed as mean ± SD from 4 independent experiments and analysed using student’s t-test.

### Heterogeneity in apoptotic priming underpins minority MOMP

We next investigated whether there was a relationship between expression of anti-apoptotic BCL-2 proteins and minority MOMP. To investigate this, we acquired super-resolution time-lapse movies of HeLa cells expressing endogenous Scarlet-BCL-2, BCL-xL or MCL-1, together with Omi-GFP and MitoTracker Deep Red. During MOMP, soluble intermembrane space proteins, including Omi, are released from mitochondria (Bock and Tait, 2020). Mitochondria retain MitoTracker Deep Red even after loss of mitochondrial integrity; thus, mitochondria that have undergone MOMP are identifiable by loss of Omi and MitoTracker retention. Surprisingly, live-cell imaging of BCL-2 family protein knock-in cells treated following treatment with ABT-737 revealed that mitochondria (determined by MitoTracker positivity) that release Omi-GFP have higher levels of BCL-2, BCL-xL or MCL-1 expression prior to MOMP (**Figure 5A-C**). Computational segmentation allowed us to distinguish BCL-2 family protein expression, which is spatially separate from the Omi signal, confirming that these mitochondria have indeed undergone minority MOMP. Quantification across a number of cells shows that mitochondria which undergo minority MOMP have increased BCL-2 family protein expression directly prior to membrane permeabilisation (**Figure 5D-F**). Furthermore, line scans revealed regions of the mitochondrial network with high BCL-2 family residency, but low Omi expression (**Supplementary Figure 4A, B**). Unexpectedly, these data reveal a correlation between increased anti-apoptotic BCL-2 expression and selective mitochondrial permeabilisation. We reasoned that this may be analogous to increased apoptotic priming at the cellular level, where high anti-apoptotic BCL-2 expression can correlate with apoptotic sensitivity in some cell types. Mitochondrial association of pro-apoptotic BAX is indicative of increasing apoptotic priming (Edlich et al., 2011; Reichenbach et al., 2017; Schellenberg et al., 2013). To investigate whether mitochondria with high BCL-2 expression may also display high BAX expression (indicative of selective, increased apoptotic priming), we generated GFP-BAX expressing BCL-2 family knock-in HeLa cells and imaged them by super-resolution microscopy. In line with the notion that mitochondria with higher BCL-2 family expression also have elevated BAX expression, we observed BAX co-localising with high BCL-2 expressing mitochondria, indicative of increased apoptotic priming (**Figure 5G-I**). These data demonstrate that inter-mitochondrial heterogeneity in anti-apoptotic BCL-2 expression and apoptotic priming underlies minority MOMP.

**Figure 5.**
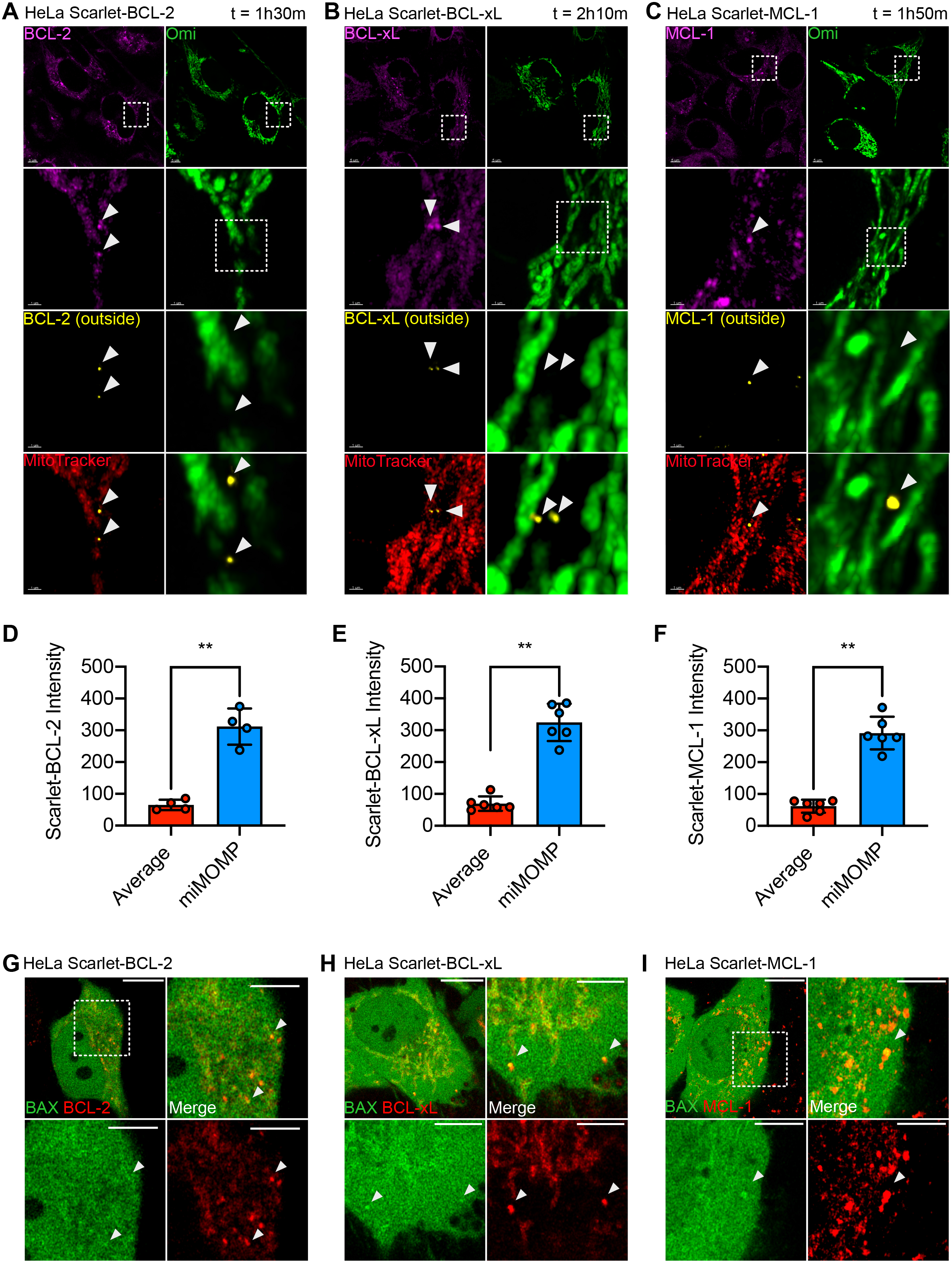
Heterogeneity in apoptotic priming underpins minority MOMP. A) Live-cell Airyscan imaging of HeLa Scarlet-BCL-2 transfected with Omi-GFP and incubated with MitoTracker Deep Red. Cells were treated with 10 µM ABT-737 for the time indicated. Images were processed with Imaris to determine BCL-2, BCL-xL or MCL-1 expression at mitochondrial areas lacking Omi-GFP expression. B) HeLa Scarlet-BCL-xL imaged as (A) C) HeLa Scarlet-MCL-1 imaged as (A) D) Quantification of Scarlet BCL-2 intensity at mitochondrial regions determined by MitoTracker Deep Red staining, but lacking Omi-GFP. Data are expressed as mean ± SD and analysed using student’s t-test. E) HeLa Scarlet-BCL-xL quantified as (D). F) HeLa Scarlet-MCL-1 quantified as (D). G) Live-cell Airyscan imaging of HeLa Scarlet-BCL-2 cells stably overexpressing GFP-BAX. Arrows indicate regions of high BCL-2 expression with high GFP-BAX expression. Scale bar = 10 µm. H) Live-cell Airyscan images of HeLa Scarlet-BCL-xL cells stably overexpressing GFP-BAX. Arrows indicate regions of high BCL-xL expression with high GFP-BAX expression. Scale bar = 10 µm. I) Live-cell Airyscan images of HeLa Scarlet-MCL-1 cells stably overexpressing GFP-BAX. Arrows indicate regions of high MCL-1 expression with high GFP-BAX expression. Scale bar = 10 µm.

### Mitochondrial dysfunction inhibits BAX retrotranslocation promoting minority MOMP

We aimed to define the underlying mechanism of mitochondrial intrinsic apoptotic priming. In healthy cells, BAX undergoes mitochondrial retrotranslocation and inhibiting this process causes BAX mitochondrial accumulation, sensitising to MOMP (Edlich et al., 2011; Schellenberg et al., 2013). Secondly, mitochondrial fusion promotes efficient oxidative phosphorylation, reducing heterogeneity in mitochondrial function (Chen et al., 2003). Given our previous data, we hypothesised that by impacting mitochondrial function, mitochondrial fission may promote BAX recruitment thereby facilitating minority MOMP. We imaged *Mfn1/2^-/-^* and *Mfn1^-/-^* MEF with MitoTracker Red, a potentiometric dye, that reports mitochondrial Δψ^m^ as a measure of mitochondrial function. Consistent with defective mitochondrial function, mitochondria in fusion defective cells (*Mfn1/2^-/-^*) displayed heterogenous MitoTracker Red signal and lower total signal than fusion competent *Mfn1^-/-^* MEF (**Figures 6A, 6B** and **6C**). We next analysed GFP-BAX localisation in *Mfn1/2^-/-^* and *Mfn1^-/-^* MEF, using fluorescence loss in photobleaching (FLIP) to help visualise mitochondrial localised GFP-BAX. Analysis of GFP-BAX localisation revealed the presence of GFP-BAX on mitochondria specifically in *Mfn1/2^-/-^* cells (**Figure 6D, Movies 2, 3**). This suggests that mitochondrial dysfunction, a consequence of defective mitochondrial dynamics, may promote GFP-BAX mitochondrial accumulation, serving as an intrinsic priming mechanism. We next asked whether mitochondrial dysfunction affected BAX localisation. HeLa cells expressing GFP-BAX and iRFP-Omp25 were treated with the uncoupler carbonyl cyanide *m*-chlorophenyl hydrazone (CCCP) to induce mitochondrial dysfunction. To facilitate visualisation of mitochondrial localised GFP-BAX, cells were treated with digitonin to selectively permeabilise the plasma membrane, as described previously (Bender et al., 2012). Inducing mitochondrial dysfunction by CCCP treatment led to robust mitochondrial recruitment of GFP-BAX (**Figure 6E, Supplemental Figure 5A, Movies 4, 5**). Immunostaining of HeLa cells with the activation specific BAX antibody 6A7 revealed BAX activation, as expected, under conditions of apoptosis (combined BH3-mimetic treatment) but not following CCCP treatment. We next measured BAX retrotranslocation rates following mitochondrial dysfunction. HeLa cells expressing GFP-BAX were treated with CCCP and retrotranslocation rates of GFP-BAX were measured by fluorescence loss in photobleaching (FLIP) (**Figures 6F and 6G, Movies 6, 7**). Re-localisation of GFP-BAX from mitochondria into the cytoplasm was reduced following CCCP treatment, demonstrating that mitochondrial dysfunction inhibits BAX retrotranslocation. These data reveal that loss of mitochondrial function, by inhibiting BAX retrotranslocation, can serve as mitochondrial intrinsic priming signal facilitating minority MOMP.

**Figure 6.**
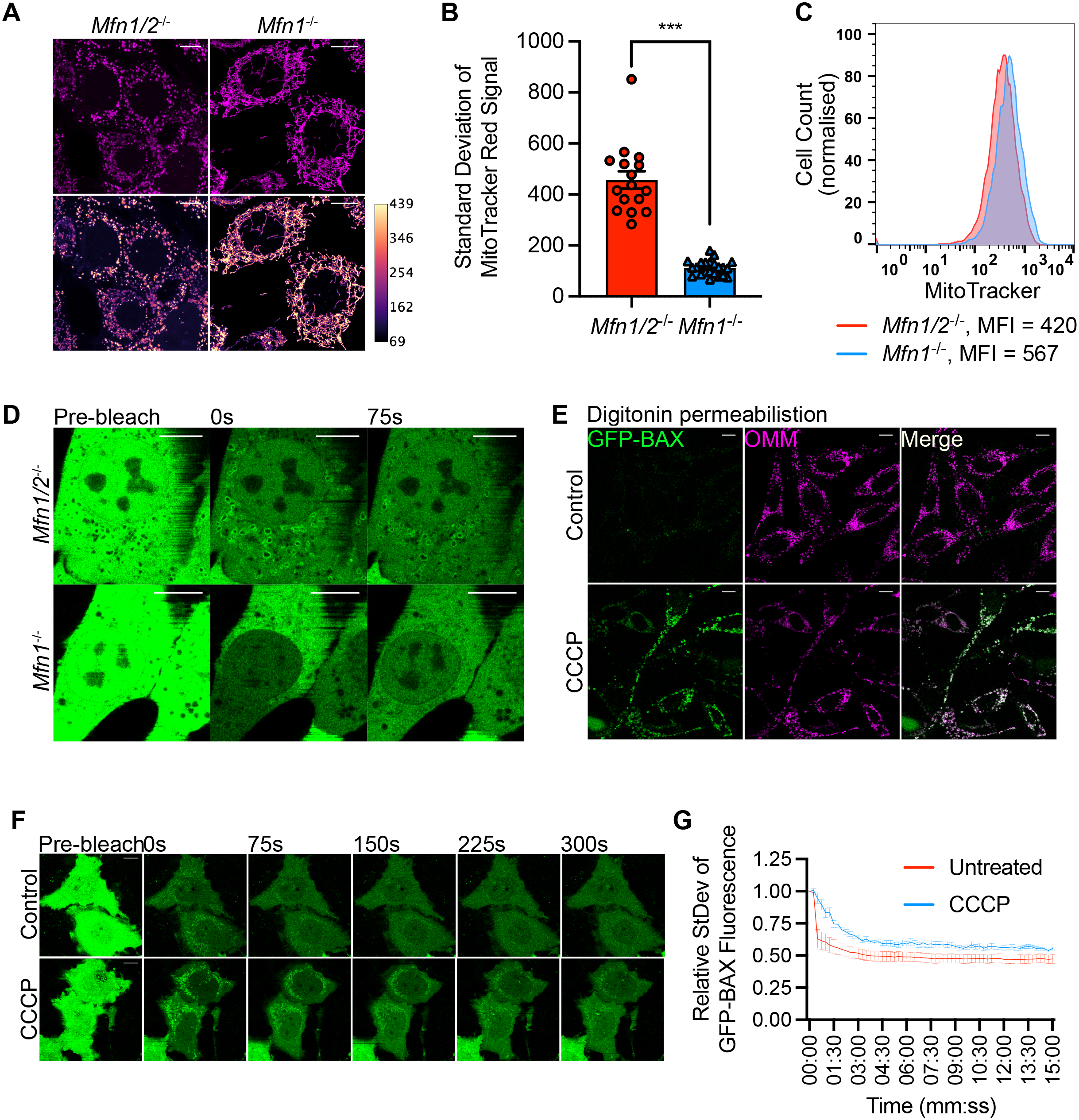
Mitochondrial dysfunction inhibits BAX retrotranslocation promoting minority MOMP. A) *Mfn1/2*^-/-^ and *Mfn1*^-/-^ MEF pulsed with MitoTracker Red and imaged. Images with magenta LUT applied are shown in lower panels. Scale bar = 10 µm. Data are representative from 3 independent experiments. B) Standard deviation of MitoTracker Red signal in *Mfn1/2*^-/-^ and *Mfn1*^-/-^ MEF pulsed with MitoTracker Red. Data are from 3 independent experiments, and analysed using student’s t-test. C) Fluorescence profiles of *Mfn1/2*^-/-^ and *Mfn1*^-/-^ MEF pulsed with MitoTracker Red. Data are representative of 2 independent experiments. D) *Mfn1/2*^-/-^ and *Mfn1*^-/-^ MEF stably expressing GFP-BAX images pre- and post-bleaching to reveal mitochondrially localised GFP-BAX. Scale bar = 10 µm. Data are representative from 3 independent experiments. See **Movie 2**. E) HeLa cells stably overexpressing GFP-BAX and iRFP-Omp25 were incubated with 20 µM digitonin to permeabilise the plasma membrane. Permeabilised cells, treated with or without 10 µM of CCCP for 30 min prior to digitonin were imaged by Airyscan microscopy. Scale bar = 10 µm. Data are representative of cells from at least 2 independent experiments. See **Movie 3**. F) HeLa cells stably expressing GFP-BAX treated with and without 10 µM of CCCP for 30 min shown pre- and post-bleaching. Scale bar = 10 µm. See **Movie 5**. G) Quantification of standard deviation of GFP-BAX signal in mitochondrial regions from cells treated with and without 10 µM CCCP from 3 independent experiments.

## Discussion

We describe that mitochondrial dysfunction, inducing mitochondrial fission, promotes DNA damage and genome instability. This process requires caspase activity, that is engaged by minority MOMP, in order to trigger DNA damage. Investigating the underlying mechanism, we find that mitochondrial dynamics affect inter-mitochondrial heterogeneity of anti-apoptotic BCL-2 expression, permitting increased apoptotic priming of fragmented mitochondria. Mitochondrial dysfunction acts as a mitochondrial intrinsic priming signal by inhibiting pro-apoptotic BAX retrotranslocation, promoting minority MOMP. Unexpectedly, by affecting mitochondrial BCL-2 heterogeneity and apoptotic priming, our data reveal crucial roles for mitochondrial dysfunction and dynamics in the regulation of minority MOMP leading to caspase dependent DNA damage and genome instability.

Our study highlights that mitochondrial dynamics are integral to minority MOMP, whereby mitochondrial fusion inhibits, and fission promotes this process. Consistent with this finding, the ability of sub-lethal apoptotic stress to engage oncogenic caspase-dependent DNA damage and genome instability was regulated in a similar manner. Moreover, we found in some cancer types, a correlation between the expression of the mitochondrial fission protein DRP1, DNA damage and mutational burden. These data support an oncogenic role for mitochondrial fission, through its capacity to promote minority MOMP and associated sub-lethal caspase activity. This also suggests that the multitude of cellular signalling pathways and stresses that impact mitochondrial dynamics, for instance as hypoxia or high glycolytic rates, might facilitate minority MOMP induced transformation (Chen and Chan, 2017; Wu et al., 2016). Indeed, we found that enforced mitochondrial fission (through MFN1/2 deletion), promoted minority MOMP induced transformation. Moreover, our study adds to the expanding interplay between mitochondrial dynamics and cancer (Chen and Chan, 2017; Gao et al., 2017; Kashatus et al., 2015; Serasinghe et al., 2015; Zhao et al., 2013).

We sought to define how mitochondrial dynamics might control minority MOMP. Surprisingly, we found extensive inter-mitochondrial heterogeneity in anti-apoptotic BCL-2 expression. This heterogeneity was supressed by mitochondrial fusion, most likely because mitochondrial fusion enables homogenous distribution of BCL-2 proteins across the mitochondrial network. As we further discuss, heterogeneity in anti-apoptotic BCL-2 expression enables differences in apoptotic priming of specific mitochondria. Interestingly, during cell death, mitochondrial variation in pro-apoptotic BAK levels have previously been found to influence the kinetics of MOMP (Weaver et al., 2014). Though myriad interconnections between mitochondrial dynamics and apoptosis exist, mitochondrial fission is largely considered a consequence of cell death. For instance, during apoptosis, extensive mitochondrial fragmentation occurs subsequent to MOMP (Bhola et al., 2009). By promoting homogenous BCL-2 expression across the mitochondrial network, our data reveal an indirect role for mitochondrial fusion in preventing minority MOMP.

We have previously found that ectopic expression of BCL-2 can lead to incomplete MOMP, consistent with BCL-2 anti-apoptotic function (Tait et al., 2010). In the current study, we find that increased expression of anti-apoptotic BCL-2 family proteins correlates with selective mitochondrial permeabilisation. While this may seem initially counter-intuitive, precedence for increased apoptotic priming, correlating with high anti-apoptotic BCL-2 levels is evident in various cancers (Certo et al., 2006; Singh et al., 2019). This is perhaps best demonstrated in high-BCL-2 expressing chronic lymphocytic leukaemia (CLL) that is often highly sensitive to the BCL-2 selective BH3-mimetic, venetoclax (Roberts et al., 2016). In healthy cells, BAX mitochondrial localisation is indicative of increased apoptotic priming (Edlich et al., 2011; Kuwana et al., 2020; Reichenbach et al., 2017; Schellenberg et al., 2013). Indeed, further investigation revealed that high pro-apoptotic BAX expression correlated with high-BCL-2 expression on intact mitochondria. Our data argue that heterogeneity in apoptotic priming exists not only between cell types, but also intracellularly, at the level of individual mitochondria.

Finally, we sought to understand how inter-mitochondrial heterogeneity in apoptotic priming might occur. Pro-apoptotic BAX is subject to constant mitochondrial retrotranslocation; inhibition of BAX retrotranslocation leads to mitochondrial accumulation, sensitising to apoptosis (Edlich et al., 2011; Schellenberg et al., 2013). We find that BAX retrotranslocation is inhibited under conditions of mitochondrial dysfunction, whereby reduction of mitochondrial inner membrane potential (Δψ_m_) promotes BAX mitochondrial localisation. Importantly, reduction of Δψ_m_, provides a mitochondrial-intrinsic signal to increase apoptotic priming. In essence, BAX retrotranslocation may serve as a barometer of cellular metabolic health. Because loss of mitochondrial function causes mitochondrial fission, it promotes minority MOMP in a two-fold manner, segregating dysfunctional mitochondria and inhibiting BAX retrotranslocation (Twig et al., 2008). Further investigation will be required to mechanistically delineate how mitochondrial function regulates BAX retrotranslocation. Moreover, we consider it likely that additional mechanisms of mitochondrial-intrinsic priming also exist.

In summary, our findings that reveal that mitochondrial dynamics regulate DNA damage and genome instability via minority MOMP induced caspase-activity. This provides a mechanism linking mitochondrial dysfunction to pro-oncogenic DNA damage. Beyond pro-tumourigenic effects, minority MOMP has also been shown to have roles in innate immunity and inflammation, as such, our findings suggest new approaches to modulate minority MOMP and its downstream functions.

## Methods

### Cell Lines

HeLa and U2OS cells were purchased from ATCC (LGC Standards). 293FT cells were purchased from Thermo Fisher Scientific.

MFN1/2^-/-^ MEF were provided by David Chan, Caltech and reconstituted with LZRS-MFN2 in our laboratory. Drp1^fl/fl^ MEF were provided by Hiromi Sesaki, Johns Hopkins University School of Medicine. All cell lines were cultured in DMEM high-glucose medium supplemented with 10% FCS, 2 mM glutamine, 1 mM sodium pyruvate, penicillin (10,000 units/ml) and streptomycin (10,000 units/ml).

To delete Drp1 from *Drp1^fl/fl^* MEF, 2 x 10^6^ cells were seeded and infected with 200 MOI high titre Ad5CMVCre (Viral Vector Core, University of Iowa) for 8 h, after which the media was replaced. Cells were used for experiments from the following day.

## METHOD DETAILS

### Generation of Scarlet-BCL-2 knock-in cell lines

We used a modified version of the knock-in strategy described in (Stewart-Ornstein and Lahav, 2016). Two vectors were used: the first vector comprises 500bp homology arm before and after the start codon of BCL-2, in between which is the Scarlet coding sequence, cloned into pUC-SP. The second vector, pSpCas9(BB)-2A-Puro (Addgene #48139) comprises Cas9 and the sgRNA targeting sequence. The following sgRNA sequences were used

Human BCL-2 5’-ATGGCGCACGCTGGGAGAAC −3’

Human BCL-xL 5-AAAAATGTCTCAGAGCAACC −3’

Human MCL-1 5’-CGGCGGCGACTGGCAATGTT −3’

To generate the knock-in cells, HeLa cells were transfected with 1 μg of homology arm vector and 1 μg of pSpCas9(BB)-2A-Puro with Lipofectamine 2000, according to the manufacturers instructions. Media was removed 5 h later, and replaced with media containing 1 μM SCR7 for 2 days. Cells were selected with 1 μg/mL puromycin for a further two days before selecting Scarlet positive clones by FACS. Cells which expressed Scarlet signal which co-localised with mitochondria we used for further experiments.

### Generation of stable overexpressing cell lines

For retroviral transduction, 293FT cells were transfected with 5 μg of plasmid, together with 1.2 μg gag/pol (Addgene #14887) and 2.4 μg VSVG (Addgene #8454) using Lipofectamine 2000. Media was changed after 6 hours and collected, filtered and used to infect cells 24 and 48 h post-transfection in the presence of 1 μg/ml Polybrene. 24 h following infection, cells were allowed to recover in fresh medium and incubated with selection antibiotic 24 h after. Cells were selected with appropriate antibiotic or FACS sorted to isolate a high-expressing population. Concentrations used for antibiotic selection were 200 μg/ml zeocin (Invivogen) or 1 μg/ml puromycin (Sigma)

For lentiviral transduction, the procdure was the same as for retroviral transduction, except 5 μg plasmid was transfected into 293FT along with 1.86 μg psPAX2 (Addgene #12260) and 1 μg VSVG (Addgene #8454) using Lipofectamine 2000.

### Generation of CRISPR knock-out cell lines

Human Drp1 and mouse Dff40 knock-out cell lines were generated by CRISPR-Cas9 gene deletion, using the lentiviral transduction protocol above. The following sequences were cloned into LentiCRISPRv2-puro (Addgene #52961)

Human Drp1: 5’- AAATCAGAGAGCTCATTCTT – 3’

Mouse Dff40: 5’- ACATGGAGCCAAGGACTCGC −3’

### Plasmids

LZRS-Drp1 was generated by cloning the Drp1 coding sequence from pcDNA3.1(+) Drp1 (Addgene #34706) into LZRS backbone using Gibson Assembly. pBABE iRFP-Omp25 was cloned by Gibson Assembly using fragments derived from pLJM2 SNAP-Omp25 (Addgene #69599) and pMito-iRFP670 (Addgene #45462). Omi-GFP (in eGFPN2) was a kind gift from Douglas Green, St. Jude Children’s Research Hospital

### Western Blotting

Cells were collected and lysed in NP-40 lysis buffer (1% NP-40, 1 mM EDTA, 150 mM NaCl, 50 mM Tris-Cl pH 7.4) supplemented with complete protease inhibitor (Roche). Protein concentration of cleared lysates was determined by Bradford assay (Bio-Rad). Equal amounts of protein lysates were subjected to electrophoresis through 10 or 12% SDS-PAGE gels and transferred onto nitrocellulose membranes, which were blocked for 1 h in 5% milk/PBS-Tween at room temperature. Membranes were incubated with primary antibody overnight at 4°C overnight. After washing, membranes were incubated with either goat-anti-rabbit Alexa Fluor 800, goat-anti-mouse Alexa Fluor 680 or goat-anti-rat DyLight 800 for 1 h at room temperature before detection using a Li-Cor Odyssey CLx (Li-Cor).

### Flow Cytometry

For measuring levels of ɣH2AX, cells were trypsinised and washed once with PBS and fixed in 4% PFA for 15 minutes at room temperature. After washing once in PBS, cells were resuspended in 300 μL and 700 μL cold ethanol added dropwise while slowly vortexing. Samples were frozen at −20°C overnight. The following day, samples were washed with PBS and blocked in 2% BSA in PBS for 1 h at room temperature and incubated with anti-ɣH2AX antibody conjugated to Alexa Fluor 647 (Biolegend) for 30 minutes protected from light. Samples were analysed on the BD LSRFortessa flow cytometer (BD Biosciences) using standard protocols.

To measure mitochondrial potential in *Mfn1/2*^-/-^ and *Mfn1*^-/-^ MEF, cells were incubated with 50nM MitoTracker CMXRos (Thermo Fisher Scientific) for 15 mins before collection. Cells were analysed on a Attune NxT flow cytometer (Thermo Fisher Scientific) using standard protocols,and analysed in FlowJo (BD).

### PALA Assay and Cad Genomic Amplification

Cells were seeded in triplicate in 6 well plates at a density of 2500 cells per well and cultured in nucleoside-free ɑ-MEM medium supplemented with 10% dialysed FBS. PALA was added at the LD_50_ dose and cells maintained until visible colonies formed. Colonies were fixed and stained in methylene blue (1% methylene blue in 50:50 methanol:water).

To assay Cad genomic amplification, DNA was extracted from PALA resistant colonies, or, in the case of control treated cells where no colonies were viable, DNA was extracted from cells passaged twenty times in DMSO, but not subjected to PALA treatment.

### Anchorage-independent growth assay

A 1% base low melting temperature agarose solution (Sigma-Aldrich) was added to 6 well plates and allowed to set. 7,500 cells were suspended in 0.6% agarose in a 1:1: ratio to achieve a final concentration of 0.3% agarose., which was added on top of base agarose. When set, the cell/agarose mix was overlaid with complete DMEM media and colonies counted 14 days later.

### q-PCR

Genomic DNA was isolated from cells using the GeneJET DNA Extraction Kit (Thermo Fisher Scientific). PCR was performed on a Bio-Rad C1000 Thermal Cycler using the following conditions: 3 min at 95°C, 40 cycles of 20 s at 95°C, 30 s at 57°C, 30 s at 72°C and a final 5 min at 72°C using Brilliant III Ultra-Fast SYBR Green qPCR Master Mix (Agilent Technologies). Relative DNA quantification was analysed by the 2^-ΔΔCt^ method. Primer sequences used are as follows:

Mouse CAD-F AAGCTCAGATCCTAGTGCTAACG

Mouse CAD-R CCGTAGTTGCCGATGAGAGG

Mouse 18S-F ATGGTAGTCGCCGTGCCTAC

Mouse 18S-R CCGGAATCGAACCCTGATT

### Microscopy

#### Fixed cell imaging

Cells were grown on coverslips and fixed in 4% PFA/PBS for 10 min, followed by permeabilization in 0.2% Triton-X-100/PBS for 15 min. Cells were blocked for 1 h in 2% BSA/PBS and incubated with primary antibodies overnight at 4°C in a humidified chamber. The following day, cells were washed in PBS and secondary antibodies added for 1 h at room temperature, before final wash steps and mounting in Vectashield antifade mounting media.

#### MOMP assay

Cells were transfected with 250ng CytoGFP and 250ng mito-mCherry for 16 h with either Lipofectamine 2000 or GeneJuice before treatment in combination with 50 nM A/C heterodimizer (Clontech). Minority MOMP was scored based on co-localisation of CytoGFP with mito-mCherry.

#### Airyscan super-resolution imaging

Super-resolution Airyscan images were acquired on a Zeiss LSM 880 with Airyscan microscope (Carl Zeiss). Data were collected using a 63 x 1.4 NA objective for the majority of experiments, although some were acquired using a 40 x 1.3 NA objective. 405nm, 561nm and 640 nm laser lines were used, in addition to a multi-line argon laser (488nm) and images acquired sequentially using the optimal resolution determined by the Zeiss ZEN software. When acquiring z-stacks, the software-recommend slice size was used. Live-cell experiments were performed in an environmental chamber at 37°C and 5% CO_2_. Airyscan processing was performed using the Airyscan processing function in the ZEN software, and to maintain clarity some images have been pseudocloured and brightness and contrast altered in FIJI (ImageJ v2.0.0).

#### Nikon A1R imaging

Confocal images were acquired on a Nikon A1R microscope (Nikon). Data were collected using a 60 x Plan Apo VC Oil DIC N2 objective. 405nm, 561nm, 638nm laser lines were used, in addition to a multi-line argon laser (488nm). Images were acquired sequentially to avoid bleedthrough. For live-cell imaging, cells were imaged in a humidified environmental chamber at 37°C and 5% CO_2_. Images were minimally processed in FIJI (ImageJ v2.0.0) to adjust brightness and contrast.

#### 3D rendering and image analysis

Z-stacks acquired on the Zeiss LSM 880 with Airyscan microscope were imported into Imaris (Bitplane, Switzerland). To segment Omi and BCL-2, a surface was created using the Omi-GFP pixel information. Masks were applied to differentiate between BCL-2 inside and outside the Omi surface. From these masks, spots were created from the BCL-2 channel and quantified based on intensity of BCL-2 on mitochondria undergoing minority MOMP.

#### Fluorescence Loss in Photobleaching

Two images were acquired before a region of interest (indicated in each figure) which overlapped the cytosol and the nucleus was bleached for 15 iterations. Photobleaching took approximately 30 seconds after which images were acquired every 15 seconds. Standard deviation of GFP-BAX signal in the cytosol was quantified using the ROI manager in FIJI and normalised.

#### Digitionin permeabilisation

Prior to digitonin permeabilisation, cells were incubated in FluoroBrite DMEM without FBS. To permeabilise the plasma membrane, 20 µM digitonin (Sigma) was added and cells imaged immediately.

#### Mitochondrial analysis

Cells stained with MitoTracker Green (100 nM) or MitoTracker Deep Red (100 nM) were imaged on the Zeiss LSM 880 with Airyscan or Nikon A1R. These images were analysed using the ImageJ plugin Mitochondrial Network Analysis (MiNA) as previously described (Valente et al., 2017). Heterogeneity of BCL-2 expression was measured by calculating the standard deviation of Scarlet and GFP signals in mitochondrial regions in FIJI (ImageJ v2.0.0).

### Live-cell viability assays

Cell viability was assayed using either an IncuCyte ZOOM or IncuCyte S3 imaging system (Sartorius). Cells were seeded overnight and drugged in the presence of 30 nM SYTOX Green (Thermo Fisher Scientific), which is a non-cell-permeable nuclear stain. Data were analysed in the IncuCyte software, and where different cell lines are compared the data are normalised to starting confluency.

### Bioinformatic Analysis

Relationship between DRP1 (*DNM1L)* expression and mutational count were investigated in TCGA PanCancer Atlas studies through cBioportal (Cerami et al., 2012; Gao et al., 2013). Studies with greater than 100 samples were analysed and samples divided into quartiles of DNM1L: mRNA expression z-scores relative to diploid samples (RNA Seq V2 RSEM). Of these, a significant association between increased mutational count in *DNM1L* mRNA highest quartile versus *DNM1L* mRNA lowest quartile was found in 2 out of 22 studies with the inverse relationship not observed in any cancer type. Mutation count in *DNM1L* quartiles was viewed in the Clinical Tab, statistical analysis of mutation count was performed by cBioportal, Wilcoxon test, q-value <0.05 was considered significant. As the relationship between DNM1L and mutational count was highly significant in Invasive Breast Carcinoma and Non-Small Cell Lung Cancer, we used these studies for further interrogation with cases of Lung adenocarcinoma selected from Non-Small Cell Lung Cancer dataset (not Lung squamous cell carcinoma). Data were downloaded from cBioportal and mutational count in *DNM1L* mRNA highest quartile versus *DNM1L* mRNA lowest quartile (mRNA expression z-scores relative to diploid samples (RNA Seq V2 RSEM)) plotted in GraphPad Prism Version 9.0.0 and statistical significance between groups calculated by Mann-Whitney test. Data points represent individual patient samples, bar is mean +SD. *DNM1L* quartiles each contain 128 samples (Lung Adenocarcinoma TCGA PanCancer Atlas dataset) or 271 samples (Breast Invasive Carcinoma TCGA PanCancer Atlas dataset). Differentially expressed proteins in DNM1L highest versus lowest quartiles were also determined in cBioportal (measured by reverse-phase protein array, Z-scores) where significant differences are determined by Student’s t-test (p value) and Benjamini-Hochberg procedure (q value). Pathway analysis was performed using gene names of proteins identified with significantly higher expression in DNM1L high versus DNM1L low quartiles (excluding phospho-specific proteins, see lists in Supplementary Table 1) in GO Biological Process 2018 through Enrichr (Chen et al., 2013; Kuleshov et al., 2016).

## Acknowledgements

Funding for this work was from Cancer Research UK Programme Foundation Award (A20145; SWGT). We thank Douglas Green (St. Jude Children’s Research Hospital), and Hiromi Sesaki (Johns Hopkins University) for reagents. We also thank Margaret O’Prey, Nikki Paul, Peter Thomason and Tom Gilbey (Beatson Institute) for excellent technical assistance, Catherine Winchester (Beatson Institute), Douglas Green and members of the Tait laboratory for critical reading of the manuscript.

## Author contributions

KC, JSR and SWGT conceived the study and designed the workplan. Experimental work: KC, JSR, CC, YE, KJC Development and contribution of reagents: KC, JSR, CC, GI. Data analysis: KC, JSR, YE, KJC, SWGT Intellectual input: KC, JSR, KJC, SWGT Manuscript writing: JSR and SWGT.

---

**Supplementary Figure 1 (related to Figure 1).**
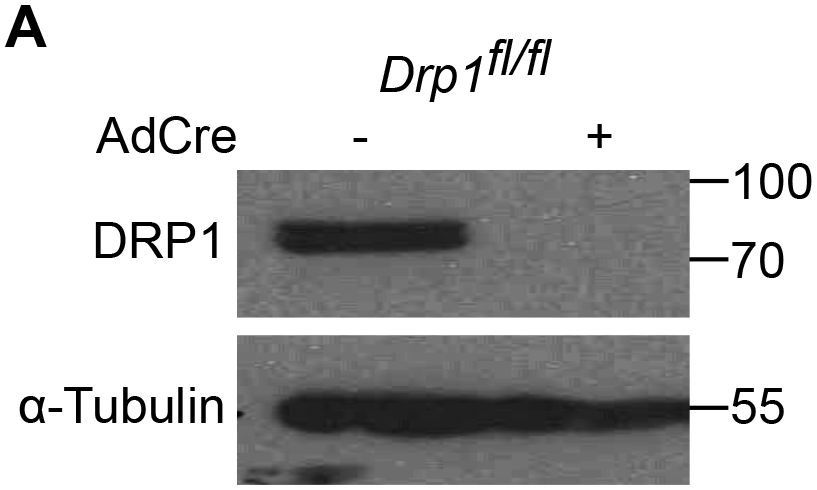
A) Western blot of *Drp1^fl/fl^* MEF with and without adenoviral Cre infection. Lysates immunoblotted for DRP1 and ɑ-tubulin.

**Supplementary Figure 2 (related to Figure 2).**
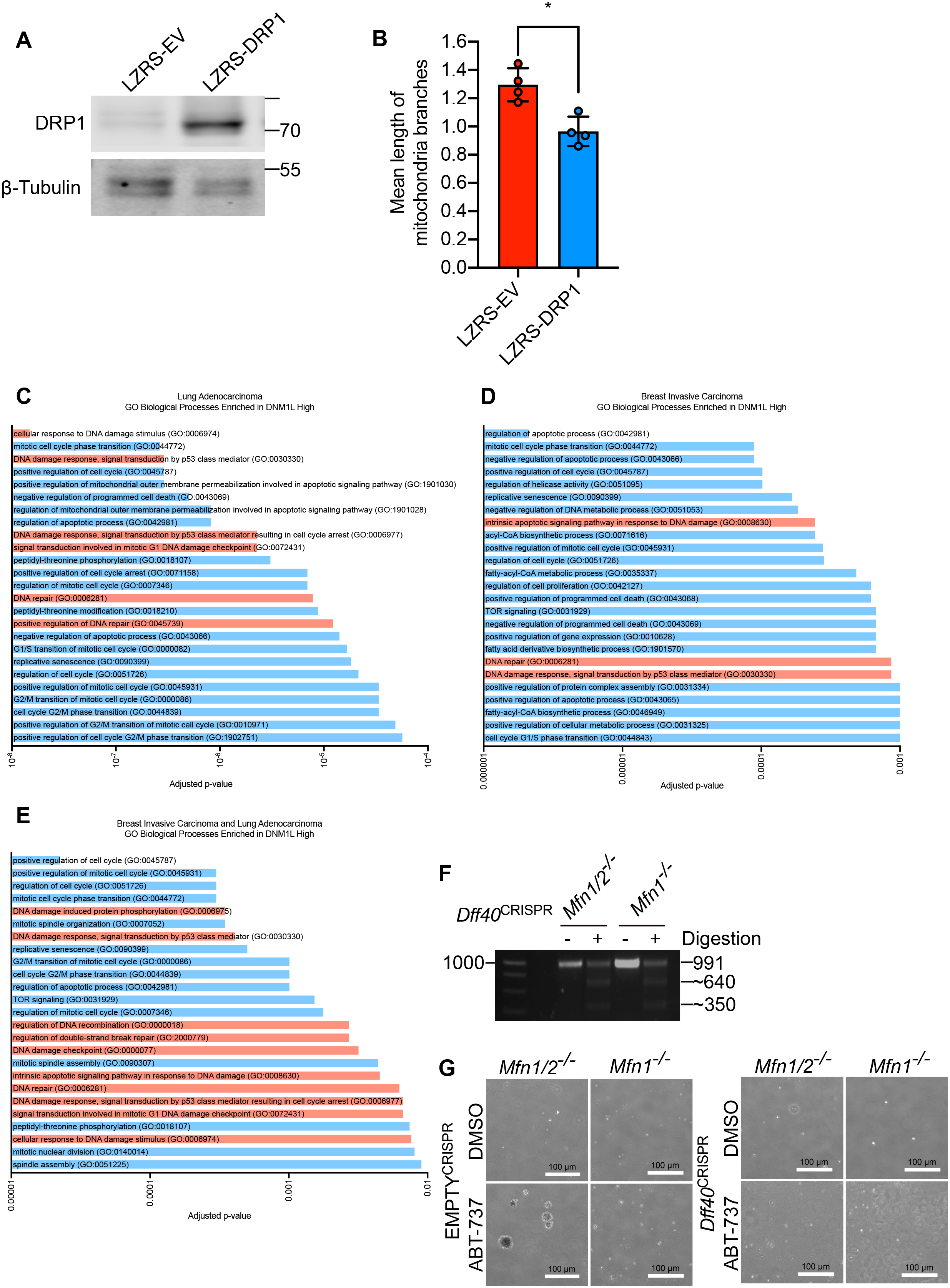
A) Western blot of MEF expressing LZRS empty control, or LZRS-DRP1. Lysates immunoblotted for DRP1 and β-tubulin. B) Quantification of mitochondrial branch size in confocal images obtained of MEF stably overexpressing LZRS empty control, or LZRS-DRP1. C) Gene ontology (GO) biological processes significantly up-regulated in DNM1L high expressing lung adenocarcinoma samples. Bars in red represent GO biological processes related to DNA damage and are expressed as adjusted p-values. D) Gene ontology (GO) biological processes significantly up-regulated in DNM1L high expressing breast invasive carcinoma samples. Bars in red represent GO biological processes related to DNA damage and are expressed as adjusted p-values. E) Gene ontology (GO) biological processes significantly up-regulated in DNM1L high expressing lung adenocarcinoma and breast invasive carcinoma samples. Bars in red represent GO biological processes related to DNA damage and are expressed as adjusted p-values. F) T7 endonuclease I mismatch assay to assay for CAD/*Dff40* deletion. G) Representative images of anchorage-independent growth of *Mfn1/2*^-/-^ and *Mfn1*^-/-^ MEF with and without CRISPR-Cas9-mediated *Dff40* deletion. Cells were passaged twenty times in 10 µM ABT-737. Scale bar = 100 µm. Data are quantified in Figure 2K.

**Supplementary Figure 3 (related to Figure 4).**
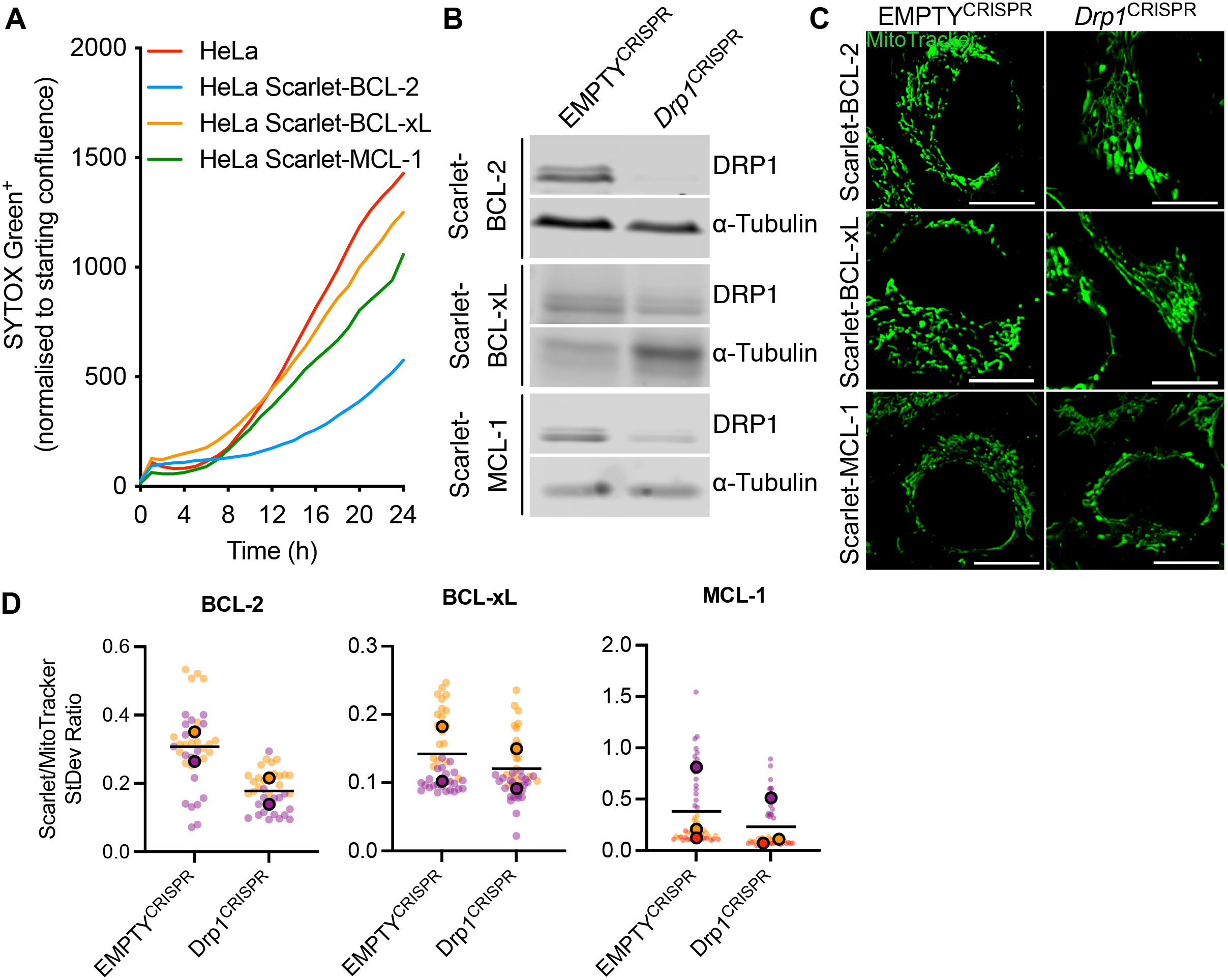
A) Live-cell IncuCyte analysis of HeLa and HeLa Scarlet-MCL-1 knockin cells treated with 10 µM ABT-737 and 2 µM S63845. Cells were incubated with SYTOX Green and SYTOX Green positivity was measured over time and normalised to starting confluency. B) Western blots of HeLa Scarlet-BCL-2, BCL-xL and MCL-1 knockin cells with CRISPR-Cas9-mediated DRP1 deletion. Lysates immunoblotted for DRP1 and ɑ-tubulin as a loading control. C) Representative Airyscan images of mitochondrial structure in HeLa Scarlet-BCL-2, BCL-xL and MCL-1 knockin cells with CRISPR-Cas9-mediated DRP1 deletion. Scale bar = 10 µm. D) Quantification of standard deviation of Scarlet to MitoTracker signal in HeLa Scarlet BCL-2, BCL-xL and MCL-1 cells, with or without CRISPR-Cas9-mediated DRP1 deletion. Data are expressed as borderless points for individual cells and points with borders represent summary data for *n* = 2 biological replicates for BCL-2 and BCL-xL, and *n* = 3 for MCL-1.

**Supplementary Figure 4 (related to Figure 5).**
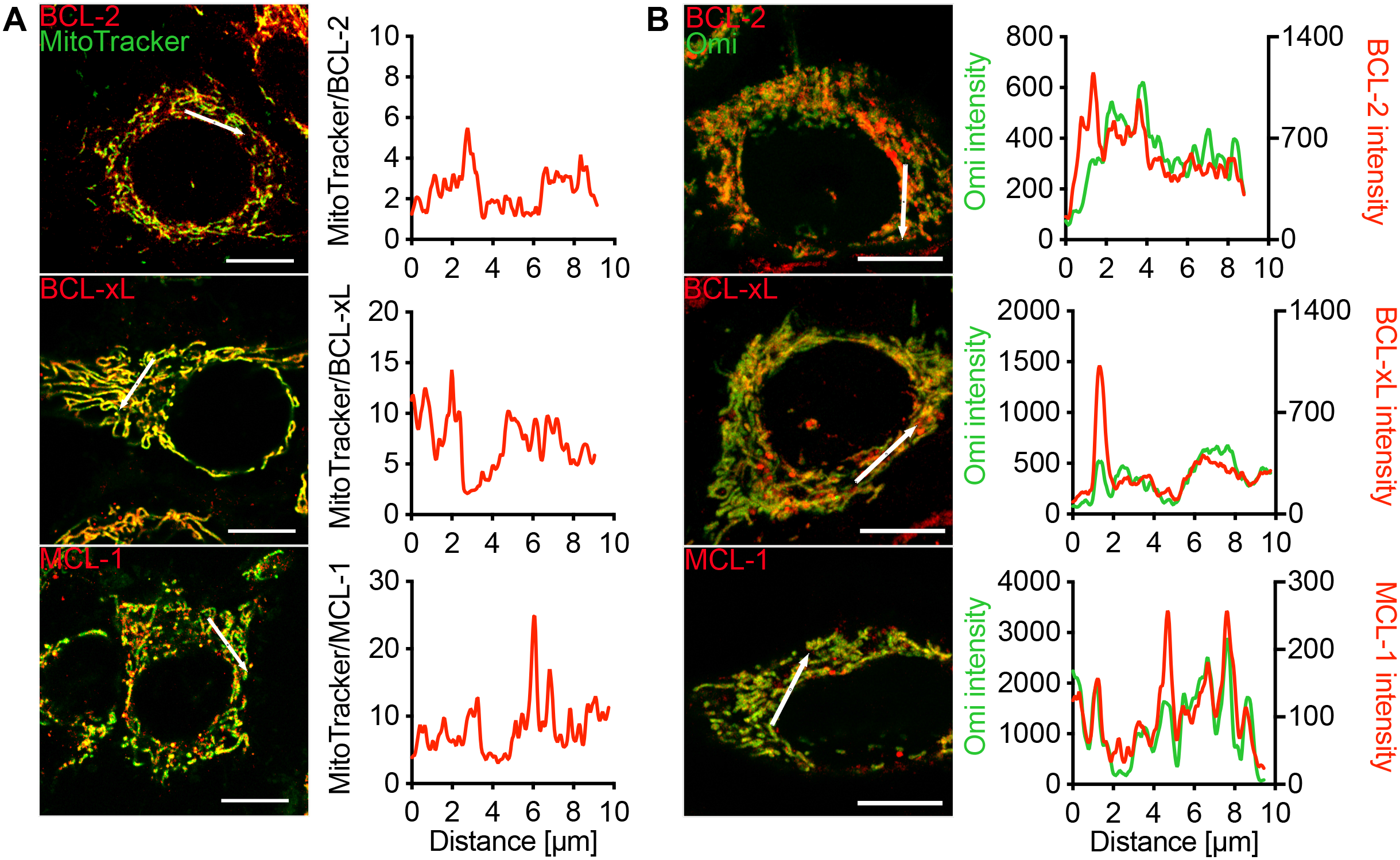
A) Airyscan images of HeLa Scarlet-BCL-2, BCL-xL and MCL-1 knockin cells and MitoTracker. Line scans show ratio of MitoTracker to Scarlet-BCL-2. Scale bar = 10 µm. B) Airyscan images of HeLa Scarlet-BCL-2, BCL-xL and MCL-1 knockin cells and Omi-GFP. Line scans show ratio of Omi-GFP intensity (green) and Scarlet-BCL-2 intensity (red). Scale bar = 10 µm.

**Supplementary Figure 5 (related to Figure 6).**
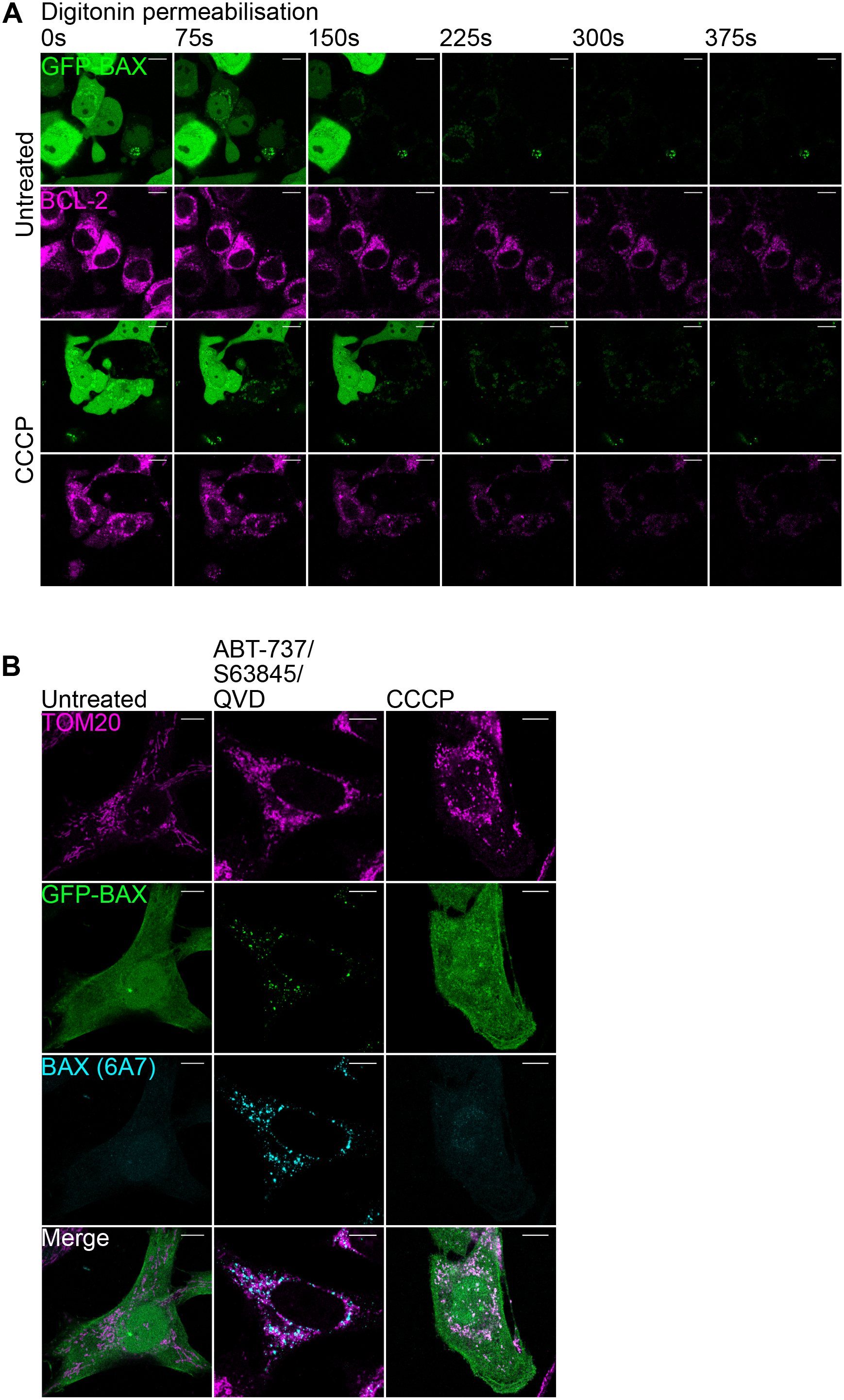
A) Images of HeLa Scarlet-BCL-2 knockin (magenta) cells stably overexpressing GFP-BAX (green) treated 10 µM CCCP for 30 min before digitonin permeabilistion. Scale bar = 10 µm. See also **Movie 4**. B) Airyscan images of fixed HeLa cells stably overexpressing GFP-BAX (green) and treated with 10 µM CCCP or 10 µM ABT-737, 2 µM S63845 and 10 µM QVD for 3 h. Cells were immunostained with anti-TOM20 (magenta) and anti-BAX 6A7 (cyan). Scale bar = 10 µm.

**Movie 1**

U2OS transfected with cytoGFP (green) and mito-mCherry (red) and treated with 10 µM ABT-737. Movie starts at 120 min. See Figure 3C.

**Movie 2**

*Mfn1/2*^-/-^ MEFs stably expressing GFP-BAX (green) and stained with MitoTracker Red (red). Cells imaged every 15 sec and bleached after 30 sec. See Figure 6D.

**Movie 3**

*Mfn1*^-/-^ MEFs stably expressing GFP-BAX (green) and stained with MitoTracker Red (red). Cells imaged every 15 sec and bleached after 30 sec. See Figure 6D.

**Movie 4**

HeLa cells stably expressing GFP-BAX (green) and iRFP-Omp25 (magenta). Cells were incubated with digitonin to permeabilise the plasma membrane and imaged every 60 sec. See Figure 6E.

**Movie 5**

HeLa cells stably expressing GFP-BAX (green) and iRFP-Omp25 (magenta) and treated with 10 µM CCCP. Cells were incubated with digitonin to permeabilise the plasma membrane and imaged every 60 sec. See Figure 6E.

**Movie 6**

*Mfn1/2*^-/-^ MEFs stably expressing GFP-BAX (green) and iRFP-Omp25 (magenta). Cells were imaged every 15 sec and bleached after 30 sec. See Figure 6F.

**Movie 7**

*Mfn1*^-/-^ MEFs stably expressing GFP-BAX (green) and iRFP-Omp25 (magenta). Cells were treated with 10 µM CCCP and imaged every 15 sec and bleached after 30 sec. See Figure 6F.

