## Supplementary Table 1 for "Mitochondrial dynamics regulate genome stability via control of caspase-dependent DNA damage"

| **Lung Adenocarcinoma** | | | | |
| --- | --- | --- | --- | --- |
| *Term* | *Overlap* | *P-value* | *Adjusted P-value* | *Genes* |
| cellular response to DNA damage stimulus (GO:0006974) | 10/329 | 1.92E-11 | 1.48E-08 | MSH6;CDKN1A;TIGAR;MRE11;BCL2L11;CCND1;CHEK2;CHEK1;CDK1;BID |
| mitotic cell cycle phase transition (GO:0044772) | 8/221 | 6.88E-10 | 2.66E-07 | CDKN1A;CCNB1;CCND1;CHEK2;EIF4EBP1;CDK1;FOXM1;EIF4E |
| DNA damage response, signal transduction by p53 class mediator (GO:0030330) | 6/82 | 1.74E-09 | 2.90E-07 | CDKN1A;CCNB1;PCNA;CHEK2;CDK1;FOXM1 |
| positive regulation of cell cycle (GO:0045787) | 6/82 | 1.74E-09 | 2.90E-07 | CCNB1;CCND1;CHEK1;EIF4EBP1;ASNS;EIF4E |
| positive regulation of mitochondrial outer membrane permeabilization involved in apoptotic signaling pathway (GO:1901030) | 5/37 | 1.88E-09 | 2.90E-07 | GSK3B;GSK3A;BCL2L11;BID;YWHAZ |
| negative regulation of programmed cell death (GO:0043069) | 9/408 | 3.91E-09 | 5.03E-07 | GSK3B;TIGAR;MRE11;CDK1;ASNS;BID;YWHAZ;BIRC2;HSPA1A |
| regulation of mitochondrial outer membrane permeabilization involved in apoptotic signaling pathway (GO:1901028) | 4/17 | 8.37E-09 | 8.20E-07 | GSK3B;GSK3A;BID;HSPA1A |
| regulation of apoptotic process (GO:0042981) | 11/815 | 8.49E-09 | 8.20E-07 | GSK3B;MRE11;BCL2L11;CHEK2;RPS6;CDK1;ASNS;BID;YWHAZ;BIRC2;HSPA1A |
| DNA damage response, signal transduction by p53 class mediator resulting in cell cycle arrest (GO:0006977) | 5/62 | 2.72E-08 | 2.28E-06 | CDKN1A;CCNB1;PCNA;CHEK2;CDK1 |
| signal transduction involved in mitotic G1 DNA damage checkpoint (GO:0072431) | 5/63 | 2.95E-08 | 2.28E-06 | CDKN1A;CCNB1;PCNA;CHEK2;CDK1 |
| peptidyl-threonine phosphorylation (GO:0018107) | 5/68 | 4.36E-08 | 3.06E-06 | GSK3B;GSK3A;CHEK2;CHEK1;CDK1 |
| positive regulation of cell cycle arrest (GO:0071158) | 5/82 | 1.13E-07 | 6.94E-06 | CDKN1A;CCNB1;PCNA;CHEK2;CDK1 |
| regulation of mitotic cell cycle (GO:0007346) | 6/165 | 1.17E-07 | 6.94E-06 | CCNB1;SMAD3;EIF4EBP1;CDK1;ASNS;EIF4E |
| DNA repair (GO:0006281) | 7/288 | 1.42E-07 | 7.83E-06 | MSH6;PCNA;MRE11;CHEK2;CHEK1;CDK1;XRCC1 |
| peptidyl-threonine modification (GO:0018210) | 5/89 | 1.70E-07 | 8.75E-06 | GSK3B;GSK3A;CHEK2;CHEK1;CDK1 |
| positive regulation of DNA repair (GO:0045739) | 4/38 | 2.54E-07 | 1.23E-05 | TIGAR;PCNA;XRCC1;FOXM1 |
| negative regulation of apoptotic process (GO:0043066) | 8/485 | 3.10E-07 | 1.41E-05 | GSK3B;MRE11;CDK1;ASNS;BID;YWHAZ;BIRC2;HSPA1A |
| G1/S transition of mitotic cell cycle (GO:0000082) | 5/105 | 3.89E-07 | 1.67E-05 | CDKN1A;PCNA;CCND1;EIF4EBP1;EIF4E |
| replicative senescence (GO:0090399) | 3/11 | 4.49E-07 | 1.82E-05 | CHEK2;CHEK1;SERPINE1 |
| regulation of cell cycle (GO:0051726) | 6/215 | 5.56E-07 | 2.15E-05 | CCNB1;CCND1;CHEK1;CDK1;FOXM1;BIRC2 |
| positive regulation of mitotic cell cycle (GO:0045931) | 4/52 | 9.19E-07 | 3.36E-05 | CCNB1;EIF4EBP1;ASNS;EIF4E |
| G2/M transition of mitotic cell cycle (GO:0000086) | 5/126 | 9.64E-07 | 3.36E-05 | CDKN1A;CCNB1;CHEK2;CDK1;FOXM1 |
| cell cycle G2/M phase transition (GO:0044839) | 5/127 | 1.00E-06 | 3.36E-05 | CDKN1A;CCNB1;CHEK2;CDK1;FOXM1 |
| positive regulation of G2/M transition of mitotic cell cycle (GO:0010971) | 3/16 | 1.52E-06 | 4.87E-05 | CCNB1;CCND1;CDK1 |
| positive regulation of cell cycle G2/M phase transition (GO:1902751) | 3/17 | 1.84E-06 | 5.68E-05 | CCNB1;CCND1;CDK1 |
| **Breat Invasive Carcinoma** | | | | |
| *Term* | *Overlap* | *P-value* | *Adjusted P-value* | *Genes* |
| regulation of apoptotic process (GO:0042981) | 13/815 | 2.36E-09 | 2.11E-06 | DIABLO;RPS6;ASNS;YWHAZ;RPS6KB1;CHEK2;KDR;CTNNB1;NF2;SQSTM1;TP53;BIRC2;TGM2 |
| mitotic cell cycle phase transition (GO:0044772) | 7/221 | 2.08E-07 | 8.89E-05 | CCNB1;CCNE2;RPS6KB1;CCNE1;CHEK2;EIF4EBP1;FOXM1 |
| negative regulation of apoptotic process (GO:0043066) | 9/485 | 2.99E-07 | 8.89E-05 | MSH2;RPS6KB1;KDR;ASNS;CTNNB1;YWHAZ;SQSTM1;TP53;BIRC2 |
| positive regulation of cell cycle (GO:0045787) | 5/82 | 5.28E-07 | 1.02E-04 | CCNB1;RPS6KB1;CHEK1;EIF4EBP1;ASNS |
| regulation of helicase activity (GO:0051095) | 3/9 | 5.71E-07 | 1.02E-04 | MSH6;MSH2;TP53 |
| replicative senescence (GO:0090399) | 3/11 | 1.12E-06 | 1.67E-04 | CHEK2;CHEK1;TP53 |
| negative regulation of DNA metabolic process (GO:0051053) | 4/43 | 1.44E-06 | 1.84E-04 | MSH6;MSH2;ERCC1;NF2 |
| intrinsic apoptotic signaling pathway in response to DNA damage (GO:0008630) | 4/48 | 2.26E-06 | 2.44E-04 | MSH6;MSH2;CHEK2;TP53 |
| acyl-CoA biosynthetic process (GO:0071616) | 3/14 | 2.46E-06 | 2.44E-04 | SCD;FASN;ACACA |
| positive regulation of mitotic cell cycle (GO:0045931) | 4/52 | 3.12E-06 | 2.79E-04 | CCNB1;RPS6KB1;EIF4EBP1;ASNS |
| regulation of cell cycle (GO:0051726) | 6/215 | 3.49E-06 | 2.83E-04 | CCNB1;CHEK1;NF2;FOXM1;BAP1;BIRC2 |
| fatty-acyl-CoA metabolic process (GO:0035337) | 3/19 | 6.50E-06 | 4.84E-04 | SCD;FASN;ACACA |
| regulation of cell proliferation (GO:0042127) | 9/740 | 9.67E-06 | 6.19E-04 | PTEN;KDR;CTNNB1;NF2;FOXM1;TP53;BAP1;BIRC2;EIF4G1 |
| positive regulation of programmed cell death (GO:0043068) | 6/257 | 9.71E-06 | 6.19E-04 | DIABLO;RPS6;CTNNB1;SQSTM1;TP53;TGM2 |
| TOR signaling (GO:0031929) | 3/23 | 1.18E-05 | 6.69E-04 | RPS6KB1;RPS6;EIF4EBP1 |
| negative regulation of programmed cell death (GO:0043069) | 7/408 | 1.22E-05 | 6.69E-04 | RPS6KB1;KDR;ASNS;YWHAZ;SQSTM1;TP53;BIRC2 |
| positive regulation of gene expression (GO:0010628) | 9/771 | 1.34E-05 | 6.69E-04 | RPS6KB1;CDH1;CCNE1;CHEK2;DVL3;CTNNB1;FOXM1;EEF2;TP53 |
| fatty acid derivative biosynthetic process (GO:1901570) | 3/24 | 1.35E-05 | 6.69E-04 | SCD;FASN;ACACA |
| DNA repair (GO:0006281) | 6/288 | 1.85E-05 | 8.64E-04 | MSH6;MSH2;CHEK2;ERCC1;CHEK1;TP53 |
| DNA damage response, signal transduction by p53 class mediator (GO:0030330) | 4/82 | 1.94E-05 | 8.64E-04 | CCNB1;CHEK2;FOXM1;TP53 |
| positive regulation of protein complex assembly (GO:0031334) | 4/88 | 2.56E-05 | 0.001041883 | ERCC1;CTNNB1;TP53;EIF4G1 |
| positive regulation of apoptotic process (GO:0043065) | 6/307 | 2.65E-05 | 0.001041883 | DIABLO;RPS6;CTNNB1;SQSTM1;TP53;TGM2 |
| fatty-acyl-CoA biosynthetic process (GO:0046949) | 3/30 | 2.68E-05 | 0.001041883 | SCD;FASN;ACACA |
| positive regulation of cellular metabolic process (GO:0031325) | 4/92 | 3.05E-05 | 0.001088903 | FASN;TP53;ACACA;EIF4G1 |
| cell cycle G1/S phase transition (GO:0044843) | 4/92 | 3.05E-05 | 0.001088903 | CCNE2;RPS6KB1;CCNE1;EIF4EBP1 |
| **Lung Adenocarcinoma and Breast Invasive Carcinoma** | | | | |
| *Term* | *Overlap* | *P-value* | *Adjusted P-value* | *Genes* |
| positive regulation of cell cycle (GO:0045787) | 4/82 | 8.47E-08 | 2.25E-05 | CCNB1;CHEK1;EIF4EBP1;ASNS |
| positive regulation of mitotic cell cycle (GO:0045931) | 3/52 | 2.70E-06 | 3.00E-04 | CCNB1;EIF4EBP1;ASNS |
| regulation of cell cycle (GO:0051726) | 4/215 | 4.04E-06 | 3.00E-04 | CCNB1;CHEK1;FOXM1;BIRC2 |
| mitotic cell cycle phase transition (GO:0044772) | 4/221 | 4.51E-06 | 3.00E-04 | CCNB1;CHEK2;EIF4EBP1;FOXM1 |
| DNA damage induced protein phosphorylation (GO:0006975) | 2/8 | 7.69E-06 | 3.48E-04 | CHEK2;CHEK1 |
| mitotic spindle organization (GO:0007052) | 3/74 | 7.85E-06 | 3.48E-04 | CCNB1;CHEK2;BIRC2 |
| DNA damage response, signal transduction by p53 class mediator (GO:0030330) | 3/82 | 1.07E-05 | 4.07E-04 | CCNB1;CHEK2;FOXM1 |
| replicative senescence (GO:0090399) | 2/11 | 1.51E-05 | 5.02E-04 | CHEK2;CHEK1 |
| G2/M transition of mitotic cell cycle (GO:0000086) | 3/126 | 3.88E-05 | 0.001010089 | CCNB1;CHEK2;FOXM1 |
| cell cycle G2/M phase transition (GO:0044839) | 3/127 | 3.98E-05 | 0.001010089 | CCNB1;CHEK2;FOXM1 |
| regulation of apoptotic process (GO:0042981) | 5/815 | 4.18E-05 | 0.001010089 | CHEK2;RPS6;ASNS;YWHAZ;BIRC2 |
| TOR signaling (GO:0031929) | 2/23 | 6.91E-05 | 0.001532567 | RPS6;EIF4EBP1 |
| regulation of mitotic cell cycle (GO:0007346) | 3/165 | 8.67E-05 | 0.001773158 | CCNB1;EIF4EBP1;ASNS |
| regulation of DNA recombination (GO:0000018) | 2/33 | 1.44E-04 | 0.0027097 | MSH6;CHEK1 |
| regulation of double-strand break repair (GO:2000779) | 2/34 | 1.53E-04 | 0.0027097 | CHEK1;FOXM1 |
| DNA damage checkpoint (GO:0000077) | 2/38 | 1.91E-04 | 0.003179542 | CHEK2;CHEK1 |
| mitotic spindle assembly (GO:0090307) | 2/46 | 2.81E-04 | 0.004395207 | CHEK2;BIRC2 |
| intrinsic apoptotic signaling pathway in response to DNA damage (GO:0008630) | 2/48 | 3.06E-04 | 0.004521309 | MSH6;CHEK2 |
| DNA repair (GO:0006281) | 3/288 | 4.48E-04 | 0.006265662 | MSH6;CHEK2;CHEK1 |
| DNA damage response, signal transduction by p53 class mediator resulting in cell cycle arrest (GO:0006977) | 2/62 | 5.11E-04 | 0.006679708 | CCNB1;CHEK2 |
| signal transduction involved in mitotic G1 DNA damage checkpoint (GO:0072431) | 2/63 | 5.27E-04 | 0.006679708 | CCNB1;CHEK2 |
| peptidyl-threonine phosphorylation (GO:0018107) | 2/68 | 6.14E-04 | 0.007425991 | CHEK2;CHEK1 |
| cellular response to DNA damage stimulus (GO:0006974) | 3/329 | 6.60E-04 | 0.007631247 | MSH6;CHEK2;CHEK1 |
| mitotic nuclear division (GO:0140014) | 2/74 | 7.27E-04 | 0.008056665 | CHEK2;BIRC2 |
| spindle assembly (GO:0051225) | 2/80 | 8.49E-04 | 0.009032491 | CHEK2;BIRC2 |
